# Splicing variants in MYRF cause partial loss of function in the retinal pigment epithelium

**DOI:** 10.1101/2025.04.21.649840

**Authors:** Gabrielle M. Rozumek, Michelle L. Brinkmeier, Bin Guan, Su Qing Wang, Catherine Tower, Nina T. Yang, Rachel Lim, Lijin Dong, D. Ford Hannum, Sayoko E. Moroi, Julia E. Richards, Robert B. Hufnagel, Lev Prasov

## Abstract

Myelin Regulatory Factor (MYRF) regulates retinal pigment epithelial (RPE) development and variants in the C-terminus are linked to isolated nanophthalmos, while loss-of-function variants cause syndromic disease. To define the molecular mechanism of this discrepancy, *in vitro* and animal studies were performed on a pathogenic C-terminal variant (p.Gly1126fs30* or dG-MYRF). ARPE-19 cells transduced with dG-MYRF revealed reduced target gene expression compared to WT-MYRF, with reduced steady state levels of C-terminal MYRF cleavage product, but intact cleavage and localization. A homozygous humanized MYRF C-terminal (*Myrf^humdG/humdG^*) mouse model was embryonic lethal by embryonic day (E) 18.5, while humanized wildtype (*Myrf^humWT/humWT^*) showed normal expression and survival. Bioinformatic analysis on integrated single cell RNA-seq from humanized E17.5 and knockout *Rx-Cre;Myrf^fl/fl^* (E15.5 and P0) mice supported shared differentially expressed genes with decreased effect size in *Myrf^humdG/humdG^* eyes. These findings, and the viability differences, support that dG-MYRF is a hypomorphic allele. Further, two novel *MYRF* splicing variants were identified in families with isolated nanophthalmos, with one confirmed to alter 40% of spliced transcripts, creating a nonfunctional isoform. These cases corroborate that isolated nanophthalmos results from hypomorphic alleles of *MYRF,* supporting a tissue-specific threshold effect and suggests that the C-terminus has unique roles in the RPE.

## INTRODUCTION

Myelin Regulatory Factor (MYRF) is a membrane associated transcription factor that was initially described for its role in oligodendrocyte maturation and myelination (1). It is a unique type II transmembrane protein that homotrimerizes in the endoplasmic reticulum (ER) lumen and auto-cleaves to release an N-terminal fragment that translocates to the nucleus and acts as a transcriptional activator (2). More recently, pathogenic variants in *MYRF* have been associated with an ocular-cardiac-urogenital syndrome featuring congenital heart defects, diaphragmatic hernias, pulmonary hypoplasia, genital abnormalities and high hyperopia (3–7). *MYRF* variants have also been described in isolated nanophthalmos, an ocular disease featuring a small but structurally sound eye, with resulting extreme farsightedness (hyperopia). C-terminal frameshift variants (impacting splicing of the final exon, and/or the amino acid sequence) have been identified in multiple large pedigrees with predominantly isolated nanophthalmos, suggesting a unique role for the *MYRF* C-terminus in the eye (8–10). However, it is not clear why some variants in *MYRF* result in isolated ocular disease and others produce syndromic phenotypes.

In the eye, MYRF is predominantly and highly expressed in the developing and mature retinal pigment epithelium (RPE) (11, 12). Conditional loss of mouse *Myrf* in the eye leads to defects in RPE development and retinal degeneration in mice, with perturbations in TGFß/BMP signaling, pigmentation, cell structure, and cell viability (12). Although the mouse models of *MYRF* do not recapitulate the eye size phenotype observed in patients, they are invaluable for understanding retinal disease pathogenesis *in vivo* (9–11).

Two large nanophthalmos families have been reported with genetic changes that lead to the same C-terminal frameshift mutation in *MYRF* causing a 31 amino acid extension in the last exon (11, 13). Interestingly, there are a cluster of variants located in the conserved C2 domain that manifest with predominantly ocular diseases (Fig. 1) (8). While the functional consequences of variants in the DNA binding and ICA domains have been well studied, there has been little to no progress on understanding the function or importance of the ER-resident, conserved C2 domain (14, 15).

**Figure 1.**
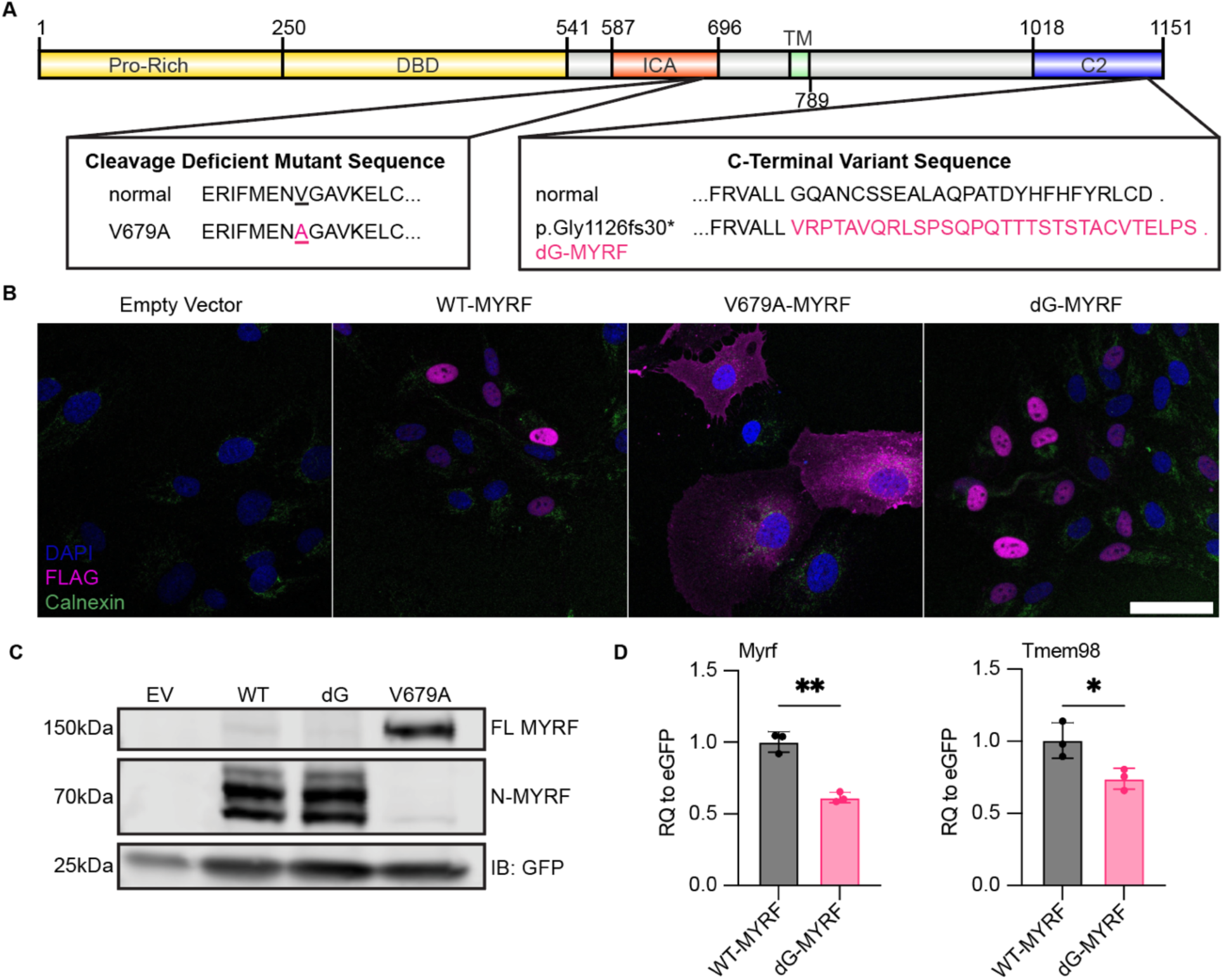
Processing of C-Terminal MYRF Variant In Vitro. (**A**) Diagram of amino acid changes in WT-MYRF, dG-MYRF, and cleavage deficient variant V679A-MYRF (ProRich: proline rich, DBD: DNA binding domain, ICA: intramolecular chaperone auto-processing, TM: transmembrane, C2: C-terminal). (**B**) Localization of FLAG tagged MYRF constructs in ARPE-19 cells show normal nuclear localization of dG-MYRF. (**C**) Western blot of transfected ARPE-19 cells shows no change in cleavage of N-terminal fragment. (**D**) qRT-PCR analysis of RNA from ARPE-19 cells transduced with dG-MYRF compared to WT-MYRF show decreased levels of transcripts for total *Myrf* and endogenous *Tmem98* mRNA (n=3). *, p<0.05, ** p<0.01, *** p<0.001

Towards that end, we generated a humanized mouse model of the C-terminal *Myrf* variant and performed *in vitro* functional assays in RPE-like cells to better understand the molecular mechanism by which C-terminal extension alleles cause disease. Our results show that the C-terminal variant retains the ability to homotrimerize, cleave, and localize properly, but has decreased C-terminal fragment steady state levels and reduced activation of downstream target gene expression. We demonstrate that the C-terminal variant acts as a reduced function allele leading to late embryonic lethality and downregulation of key pathways involved in RPE development. Further, we identified two additional rare splice site variants in *MYRF* in families with isolated nanophthalmos supporting this as a more common mechanism for disease pathogenesis. Together, our studies support that partial loss of function alleles in *MYRF* contribute to isolated nanophthalmos and provide mechanistic insights into the role of the MYRF C2 domain.

## RESULTS

### C-Terminal MYRF Variant is Processed Normally but Shows Decreased Stability and Transcriptional Activity

C-terminal *MYRF* variants and splicing variants are associated with predominantly ocular phenotypes and have familial inheritance, while early truncating and ICA and DBD variants are associated with syndromic phenotypes and predominantly occur *de novo* (8, 13, 15–17). Given this discrepancy, the C-terminal frameshift alleles could either be acting as dominant negative or hypomorphic alleles. To distinguish these models, we evaluated localization, cleavage, and protein stability of the C-terminal variant *in vitro* in RPE-like ARPE-19 cells. When overexpressed with lentiviral vectors, both Wild-type (WT) MYRF and C-terminal frameshift MYRF (dG-MYRF) showed nuclear localization, while a known cleavage deficient mutant V679A-MYRF (14) displayed only cytoplasmic localization and was excluded from the nucleus (Figure 1A-B). As homotrimerization at the ER lumen is necessary for cleavage of the N-terminal MYRF fragment and its subsequent translocation to the nucleus (1, 2, 18, 19), our results support the dG-MYRF retains ability to homotrimerize and cleave.

To further evaluate this, we used Western blotting to systematically define the cleavage dynamics (ratio of full length to cleaved) of the dG-MYRF variant compared to WT-MYRF in ARPE-19 cells. We observed faint bands for full length FLAG-MYRF (160kDa) band in both WT-MYRF and dG-MYRF and strong N-MYRF cleavage products (~70kDa). In contrast, the cleavage deficient mutant, V679A-MYRF, shows a singular strong full length MYRF band (Figure 1C). Given normal cleavage and localization, we next evaluated protein stability through computational tools examining degree of disorder <u>P</u>redictor <u>o</u>f <u>N</u>atural <u>D</u>isordered <u>R</u>egions (PONDR) and protein folding (AlphaFold2) (20–23). PONDR uses neural networks trained on ordered and disordered regions of short and/or long regions of amino acid sequences from NMR or x-ray crystallography data to predict intrinsic disordered regions. Using PONDR, dG-MYRF was predicted to alter the C-terminus from a highly ordered to highly disordered structure (Supplemental Figure 1A). *De novo* structural modeling using AlphaFold2 predicted the loss of a ß-sheet and alpha-helix structure in the conserved C2 domain of the C-terminal frameshift variant of *MYRF* (Supplemental Figure 1, B and C). To experimentally validate this, cycloheximide pulse-chase experiments were conducted in ARPE-19 cells transduced with WT-MYRF and dG-MYRF. Protein extracts were analyzed by Western blotting using antibodies to the N-terminus or C-terminus of MYRF. The rates of decay of the C-terminal and N-terminal MYRF cleavage product were unchanged in the dG-MYRF variant following a 24 hour chase (Supplemental Figure 2A). However, steady state levels of C-terminal cleavage product for dG-MYRF were reduced compared to wild-type (Supplemental Figure 2, B and C). This was unlikely due to reduced epitope detection because the antibody was raised to an epitope present in both forms of MYRF (anti-MYRF^393-766AA^) and the C2 domain is predicted to fold independently (1).

To evaluate the functional impact of the dG-MYRF variant on transcriptional activity, we tested the ability of dG-MYRF to autoregulate *MYRF* transcription and its downstream target *TMEM98* using qRT-PCR, as compared to an internal transduction GFP control, present in each of the constructs. ARPE-19 cells transduced with dG-MYRF showed decreased levels of endogenous *MYRF* and *TMEM98* mRNA expression (relative to GFP) as compared to WT-MYRF (Figure 1D). Thus, the dG-MYRF variant does not impact cleavage dynamics or nuclear localization, but it reduces transcriptional activity. There is also reduction in the steady state level of C-terminal cleavage product, possibly through altered folding of the C-terminal C2 domain of *MYRF*.

### Mouse Model of Human C-Terminal MYRF is Prenatal Lethal

To understand the pathogenesis of nanophthalmos in patients with the *MYRF* c.3376-1G>A variant, we used CRISPR/Cas9 homology directed repair to generate a humanized mouse model that mimics the frameshift variant detected in patients, as well as a matched humanized wild-type control (Figure 2A). In this model, we replaced mouse exon 26 with either the human wildtype or human C-terminal frameshift exon 27 DNA sequence, including the endogenous 3’UTR sequence, and fused this with mouse exon 25 (Supplemental Figure 3). We previously showed that *Myrf* is highly expressed in RPE via qRT-PCR on RNA from optic cups and RNAscope *in situ* hybridization (11, 12). To confirm proper cell specificity and expression of the humanized alleles of *Myrf*, we used a validated *Myrf* RNAscope probe to evaluate *Myrf* expression in *Myrf^humdG/humdG^*and *Myrf^humWT/humWT^* eyes at E15.5. These studies showed consistent *Myrf* expression pattern and levels among *Myrf^humWT/humWT^*and *Myrf^humdG/humdG^* eyes within the RPE (Figure 2B). To further define the function of the dG allele, we first evaluated survival of *Myrf^humdG/humdG^* embryos. We observed that homozygous *Myrf^humdG/humdG^* embryos were never detected after birth, while *Myrf^humWT/humWT^* mice were viable and fertile. *Myrf^humdG/humdG^* embryos were present in normal Mendelian ratios at E12.5-17.5 but underrepresented at E18.5 (chi^2^ prob = 0.015) and at birth (P0) (chi^2^ prob = 0.042) (Figure 2C).

**Figure 2.**
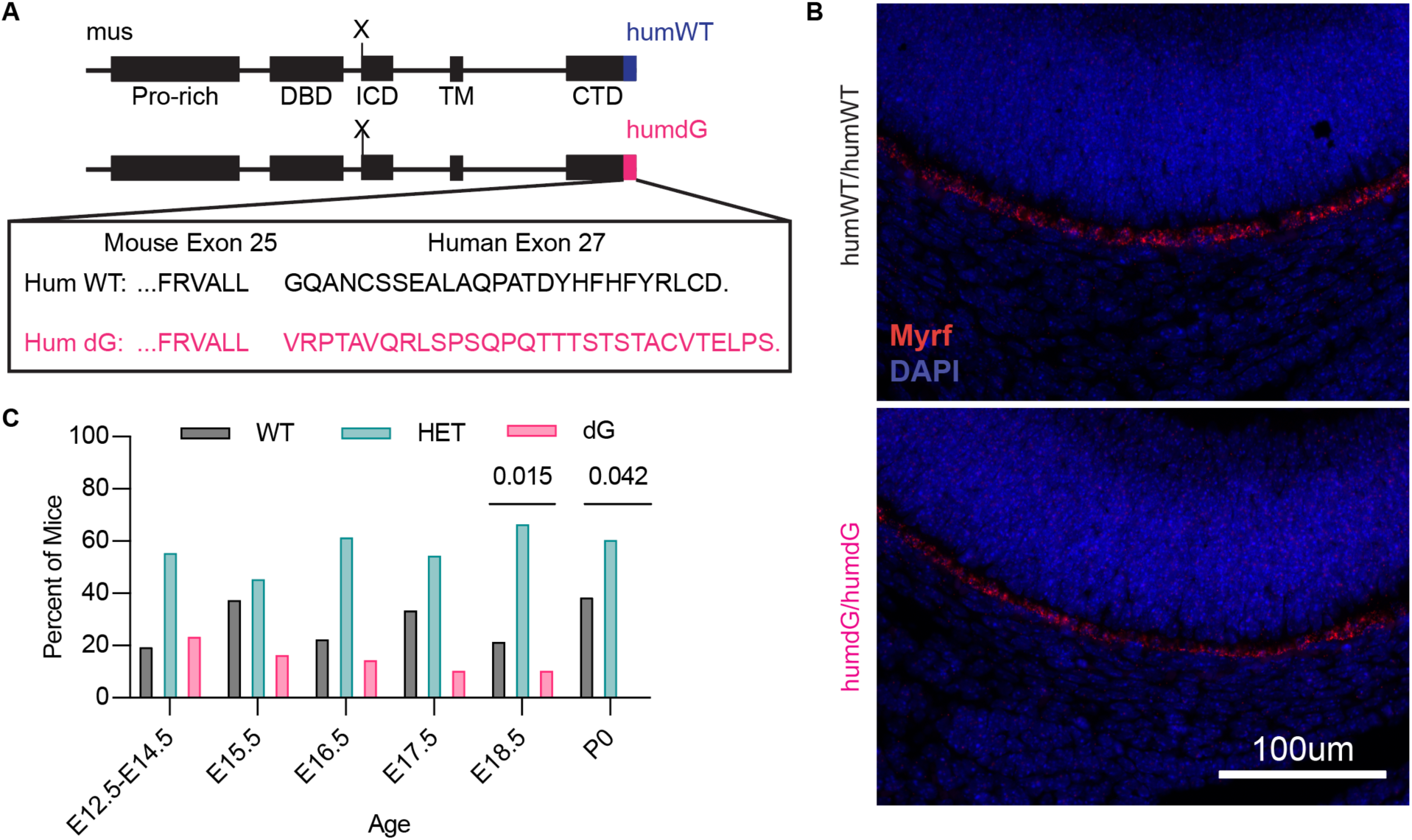
Homozygous *Myrf humdG* mice are embryonic lethal. (**A**) Schematic showing the (humdG) fusion allele amino acid sequence changes in the humanized mouse model. (**B**) In situ hybridization using RNAScope detected *Myrf* mRNA in humanized humWT and humdG mouse RPE (n=4-5 per genotype), scale bar = 100µm. (**C**) Progeny from intercrossing *Myrf^humWT/humdG^*mice exhibit skewed Mendelian ratios at E18.5 (chi^2^ p = 0.015) and P0 (p=0.04).

Analysis of ocular histology in E16.5 homozygous *Myrf^humdG/humdG^*mice revealed normal pigmentation, which differed from the severe and early onset depigmentation phenotype we observed in our *Rx>Cre Myrf^fl/fl^*mice (Supplemental Figure 4) (11). *Tmem98*, a direct target of MYRF, is expressed at similar levels in control eyes (*Myrf^humWT/humWT^*^)^ and *Myrf^humdG/humdG^* eyes at the RNA and protein level (Supplemental Figure 5) (11, 24). We cannot rule out the possibility that there are small expression differences not detected by the mRNA and immunostaining assays. However, the *Rx>Cre Myrf^fl/fl^* eyes had profoundly reduced expression of *Tmem98* (*12*). The lethality of *Myrf^humdG/humdG^* mice confirm the pathogenicity of the C-terminal *Myrf* mutant allele, and the lack of obvious ocular defects suggest that it likely functions as a hypomorphic (partial loss of function allele) rather than null allele.

### *Myrf^humdG/+^* have Normal Eye Size, Retinal and RPE Morphology, and Retinal Function

The *Rx>Cre Myrf^fl/+^* mice had white spots in the retina and *Rx>Cre Myrf^fl/fl^* mice had severe retinal degeneration, but no eye size phenotype was observed in either genotype (11). We systematically profiled *Myrf^humWT/+^* and *Myrf^humdG/+^* mice over the course of 1 year to determine whether changes ocular dimensions, retinal structure, or function would emerge over time. Non-invasive imaging (SD-OCT and fundus photography) and electroretinography (ERG) were carried out on the same mice over the course of 1 year at 3-month intervals. After 1 year, *Myrf^humdG/+^* show no significant different in eye size, axial length, corneal diameter, anterior chamber depth, or retinal thickness, and only a modest decline in vitreous chamber depth (0.502± 0.197, p = 0.0342) compared to controls (Figure 3A-G). Additionally, scotopic and photopic ERGs showed no decrease in visual function consistent with the SD-OCT findings of normal retinal thickness and morphology (Figure 3 H-J). Fundus photography showed little to no signs of atrophy after 1 year (Figure 3B-C). To investigate for more subtle changes to RPE morphology similar to those that were evident in *Rx>Cre Myrf^fl/fl^* eyes (11), RPE flat mounts were stained with phalloidin, segmented using REShAPE software (25), and evaluated for morphometric features (Supplemental Figure 6). These analyses revealed no differences in the median cell area (p = 0.899), aspect ratio (p = 0.847), or hexagonality (p= 0.7568) of 1 year old *Myrf^humdG/+^*eyes compared to *Myrf^humWT/+^* controls. There was also no change in the median number of neighboring RPE cells between genotypes. These lack of RPE pathology supports the idea that the *Myrf^humdG^* allele is hypomorphic.

**Figure 3.**
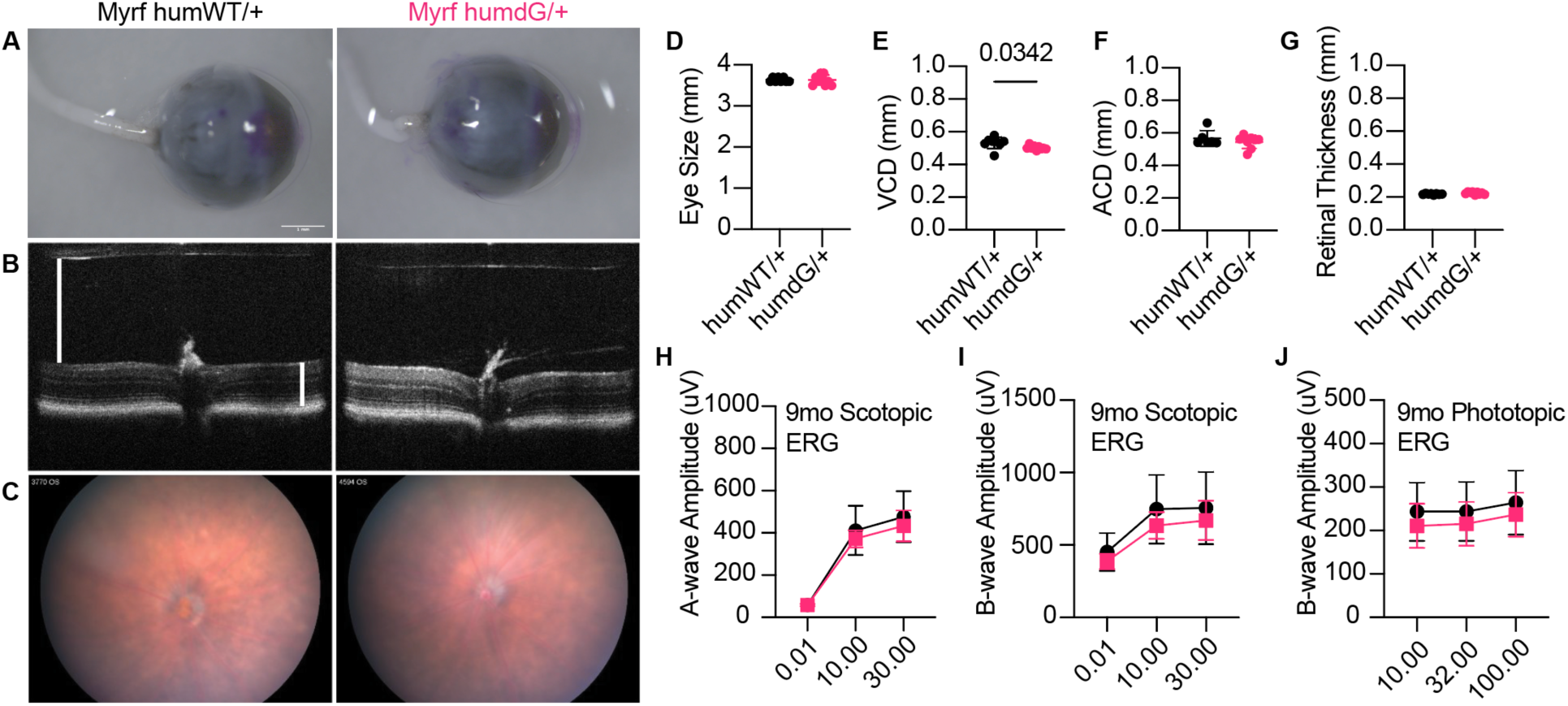
Heterozygous *Myrf^humdG/+^*mice show no gross morphological or retinal eye phenotype after 1 year. (**A**) Images of enucleated eyes showing similar eye size. Scale bar = 1mm. (**B**) Spectral domain optical coherence tomography (SD-OCT) images showing no change in overall thickness or individual retinal layers with white lines representing measurements for vitreous chamber depth (VCD) and retinal thickness. (**C**) Fundus photographs showing no signs retinal white spots in *Myrf^humdG/+^* compared to controls. (**D-H**) Quantitative assessment of eye size (D), VCD (E), anterior chamber depth (F), and retinal thickness (G). (**I-K**) Retinal function measured by scotopic electroretinogram (ERG) (H,I) and photopic ERG (J) showed no differences (n=6-8 per genotype).

### Single Cell RNA-sequencing Shows Altered RPE Gene Regulatory Networks in *Myrf^humdG/humdG^* Eyes

To better understand the how the C-terminal MYRF variant impacts eye development we performed single cell RNA-sequencing on E17.5 optic cups of *Myrf^humWT/humWT^*and *Myrf^humdG/humdG^* mice using the 10X genomics platform (Figure 4A). We chose E17.5 because it is slightly after the initiation of RPE pigmentation at E15.5 and just before Mendelian ratios became skewed at E18.5. We collected 11,281 and 11,322 cells for *Myrf^humWT/humWT^* and *Myrf^humdG/humdG^*, respectively. The median genes per cell were comparable in wild-type (2429) and variant (2415) samples. Quality control filtering was performed to filter out of dead cells, doublets, and poor-quality cells (12). After integration with previously published datasets from conditional knockout Myrf eyes (Rx>Cre Myrf^fl/fl^ and Myrf^fl/fl^) at E15.5 and P0 we performed unsupervised clustering and were able to identify 19 clusters including all the major cell types within the optic cup using previously established cell type specific markers (Supplemental Figure 7) (12). All clusters were present in both the *Myrf^humWT/humWT^* and *Myrf^humdG/humdG^*mice (Figure 4B), but there was a slight reduction in the RPE cell proportions in *Myrf^humdG/humdG^* mice (5.0%) compared to control (6.4%) (Supplemental Figure 7C-D).

**Figure 4.**
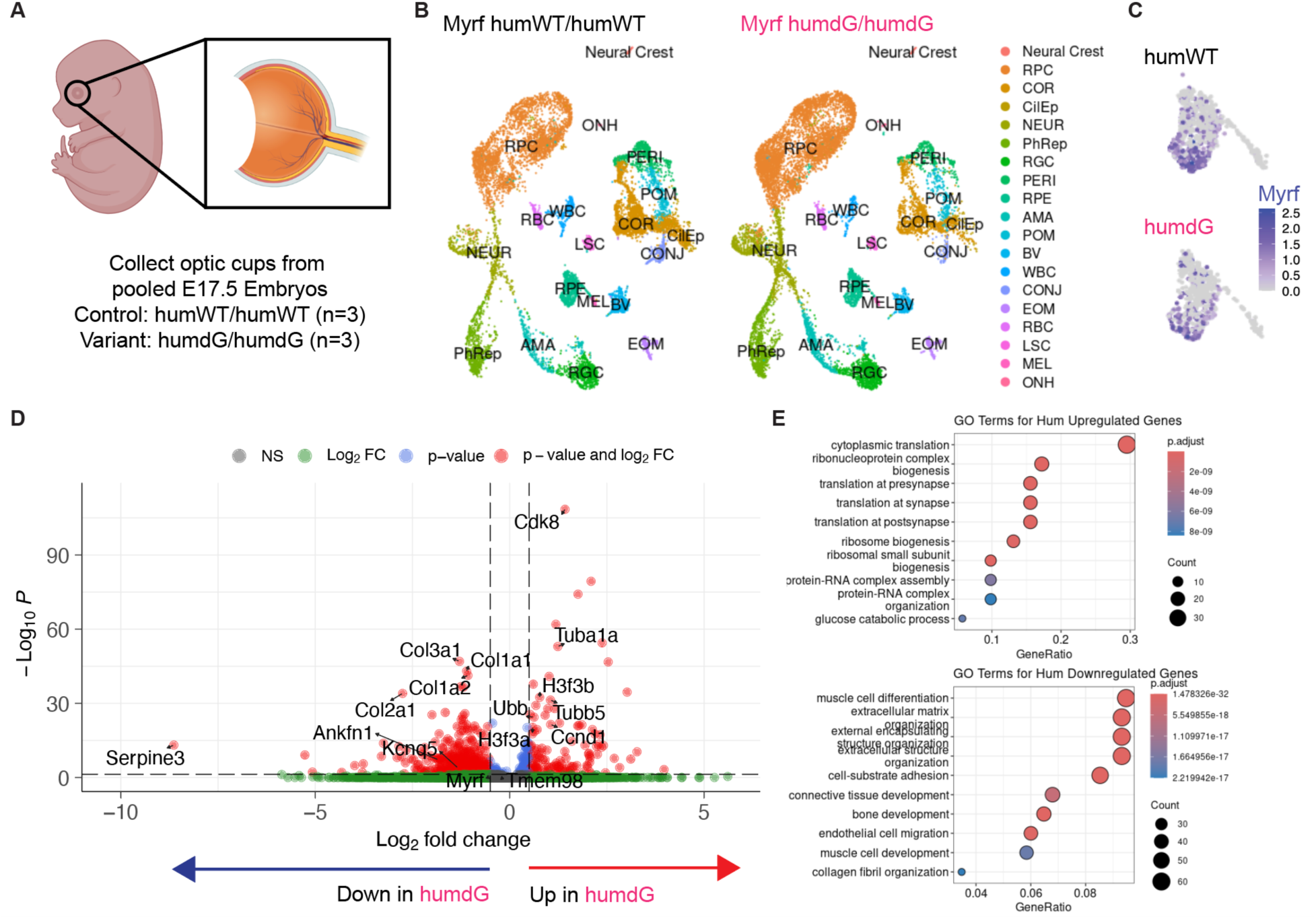
Single cell RNA-seq reveals molecular changes in RPE of mice carrying C-Terminal *Myrf* Allele (M*yrf^humdG/humdG^*). **(A)** Schematic depicting optic cups and samples used for scRNA-seq including control and variant mouse optic cups (n=3 pooled per genotype). (B) UMAPs of control vs. *Myrf* C-terminal variant mouse optic cups demonstrating cell type distributions. (C) Feature plot demonstrating expression of *Myrf* mRNA in both control and variant mice within the RPE cluster. (D) Volcano plot of differentially expressed genes in *Myrf^humdG/humdG^* mice compared to control (Myrf^humWT/humWT^). (E) Top ten Gene Ontology (GO) Biological Pathway terms for upregulated and downregulated genes.

*Myrf* expression was detected exclusively in the RPE cluster (Figure 4C), consistent with previous results. For that reason, we focused our downstream analysis on the RPE cluster. We performed differential gene expression analysis between *Myrf^humWT/humWT^*and *Myrf^humdG/humdG^* RPE clusters. We found 735 upregulated and 170 downregulated differentially expressed genes (DEGs) with avg_log2FC > 0.25 and adjusted p-value <= 0.1 (Figure 4D), Myrf^humdG/humdG^ relative to Myrf^humWT/humWT^ controls. Downregulated genes included extracellular matrix related (ECM) genes C*ol3a1*, *Col2a1*, *Col1a1*, *Col1a2* produced by the RPE and play an important role in structure and barrier maintenance (26). Interestingly, some of the most downregulated genes in the variant mice are related to eye size disorders including *Serpine3* (avglog2FC = −1.11, padj = 2.76E-14), *Ankfn1* (avglog2FC = −1.138, padj = 2.72E-09), and *Kcnq5* (avglog2FC = −0.81, padj=6.39E-05) (27–31). Upregulated genes included cell cycle genes *Ccnd1* and *Cdk8*, histone variant components *H3f3b* (avglog2FC = 0.71, padj = 3.69E-34) and *H3f3a* (avglog2FC = 0.56, padj = 3.30E-23), and ubiquitin protein ligase binding genes *Tubb5, Ubb* (avglog2FC = 0.59, padj = 2.32E-27), *Tuba1a* (avglog2FC = 0.32, padj = 0.03). We used overrepresentation analysis (ORA) to identify the top 10 gene ontology terms for Biological Processes and Molecular Function pathways in humanized variant mice (*Myrf^humdG/humdG^*) and conditional knockout (*Rx-Cre;Myrf^fl/fl^*) relative to their matched controls (Myrf^humWT/humWT^ and Myrf^fl/fl^). We looked for overlap in top GO terms enriched in Myrf^humdG/humdG^ and condition knockout compared to controls and found shared downregulation of extracellular matrix organization (GO:0030198) and extracellular structural organization (GO:0043062). In addition, shared upregulated pathways included structural constituent of ribosome (GO:0003735), structural molecule activity (GO:0005198), rRNA binding (GO:0019843), and translation (GO:0006412) (Figure 4E). In addition, GO analysis identified many upregulated pathways in both humanized and knockout mice related to metabolism, mitochondrial, and ribosomal function. Thus, the humanized C-terminal variant mutant (*Myrf^humdG/humdG^*) effects the RPE transcriptome in a manner like the conditional knockout (*Rx-Cre;Myrf^fl/fl^*), leading to the downregulation of key genes involved in RPE maintenance and development.

We extended the differential gene expression analysis of the humanized C-terminal variant allele (*Myrf^humdG/humdG^*) and conditional knockout (*Rx-Cre;Myrf^fl/fl^*) by examining individual genes that were affected, including the direction and magnitude of the effect. After integration, we performed differential gene expression analysis on these two conditions and looked for genes that had an avg_log2FC >= abs(0.25) and adjusted p-value <= 0.1 in both the C-terminal variant and conditional knockout mice compared to controls. We found a total of 55 shared DEGs, with 27 downregulated and 28 upregulated genes respectively (Figure 5A, Supplemental Table 1). Key genes important for RPE development and maintenance were concordant but the magnitude of the change was less in *Myrf^humdG/humdG^*compared to *Rx>Cre Myrf^fl/fl^* mice (Supplemental Figure 8). Some of the concordant downregulated genes and pathways included ECM matrix (*Upk3b*, *Upk1b*, *Cdh3*), Wnt signaling (*Tcf4*), TGFß inhibition (*Wfkkin2*), and *Myrf*. The concordance of upregulated genes included increases TGF-Beta signaling (*Id1* and *Id3*) and glucose metabolism (*Gapdh, Ldha, Eno1*) (Figure 5B). Gene set enrichment analysis (GSEA) was performed on separately on both *Myrf^humdG/humdG^* and *RxCre;Myrf^fl/fl^* paired with matched controls, Myrf^humWT/humWT^ and Myrf^fl/fl^ respectively. GSEA results from *Myrf^humdG/humdG^* RPE emphasized downregulation of extracellular matrix and bone development/differentiation pathways consistent with our prior reports which showed similar enrichment in *Rx>Cre Myrf^fl/fl^* RPE. Notably, pathways related to eye and sensory organ development were absent in the top GSEA results for *Myrf^humdG/humdG^*, while they dominated the top results for *Rx>Cre Myrf^fl/fl^*. (Figure 5, C and D). Finally, treeplots were generated to show hierarchical clustering of enriched GO biological process terms. These plots highlighted shared (eye and visual system development) and unique pathways (cell adhesion and GTPase signaling) and defined connective tissue, adhesion, and chondrocyte as high-frequency words in the C-terminal *Myrf* variant as compared to the conditional knockout of *Myrf* (Figure 5, E and F). Together, these data support shared pathogenic mechanisms between the knockout model and the C-terminal variant at the level of RPE and a hypomorphic effect.

**Figure 5.**
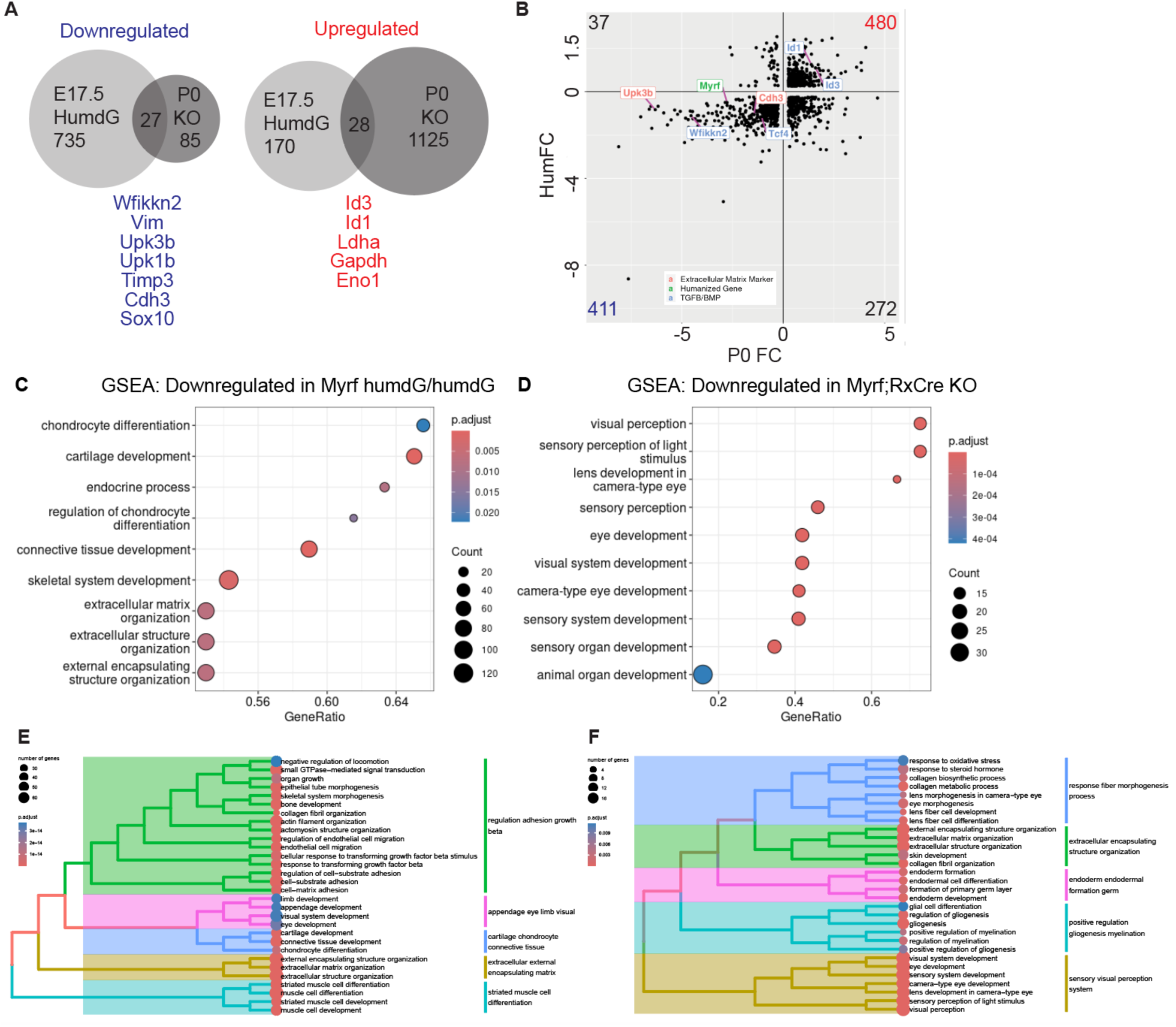
*Myrf^humdG^*C-Terminal Variant Acts as a Hypomorphic Allele. **(A)** Comparison of DEGs in RPE cluster of E17.5 humanized C-terminal variant (*Myrf^humdG/humdG^*) vs. conditional knockout (*Rx>Cre Myrf^fl/fl^*) mice with key shared genes highlighted. (B) Plot of Myrf^humdG/humdG^ vs. RxCre;Myrf^fl/fl^ fold change in gene expression for genes expressed in both datasets (log2FC > 0.25, adjusted p-value < 0.1) showing strong concordance of up and downregulated DEGs between the humanized allele and conditional knockout. (C, D) Gene set enrichment analysis (GSEA) showing top ten downregulated pathways in *Myrf^humdG/humdG^*vs. *Rx>Cre Myrf^fl/fl^* mice. (E-F) Treeplots showing hierarchical clustering of enriched GO biological process terms in *Myrf^humdG/humdG^* (E) vs. *Rx>Cre Myrf^fl/f^*^l^ mice (F)

### Novel Deep Intronic Variants in MYRF in Nanophthalmos Patients Alter Splicing

As our in vitro and transcriptomic findings suggest a hypomorphic effect for the dG-MYRF variant, we reasoned that other *MYRF* splicing variants may be more likely to cause ocular only phenotypes, either by affecting only the C-terminal domain exons or by causing a partial reduction in functional RNA or protein products. Towards that end, we used short-read, panel-based or whole genome sequencing (WGS) on a cohort of 89 families with nanophthalmos/high hyperopia that did not have identified genetic diagnoses (32). From this cohort, two families were identified with deep intronic variants in *MYRF* (NM_001127392.3 MYRF c.460+167G>A in P01965 from family 1 and c.3194+122A>G in P04818 and P04825 from family 2) (Figure 6A-B). No other rare deleterious genetic variants in nanophthalmos-associated genes (*MYRF*, *PRSS56*, *MFRP*, *CRB1*, BEST1) were identified in either family.

**Figure 6.**
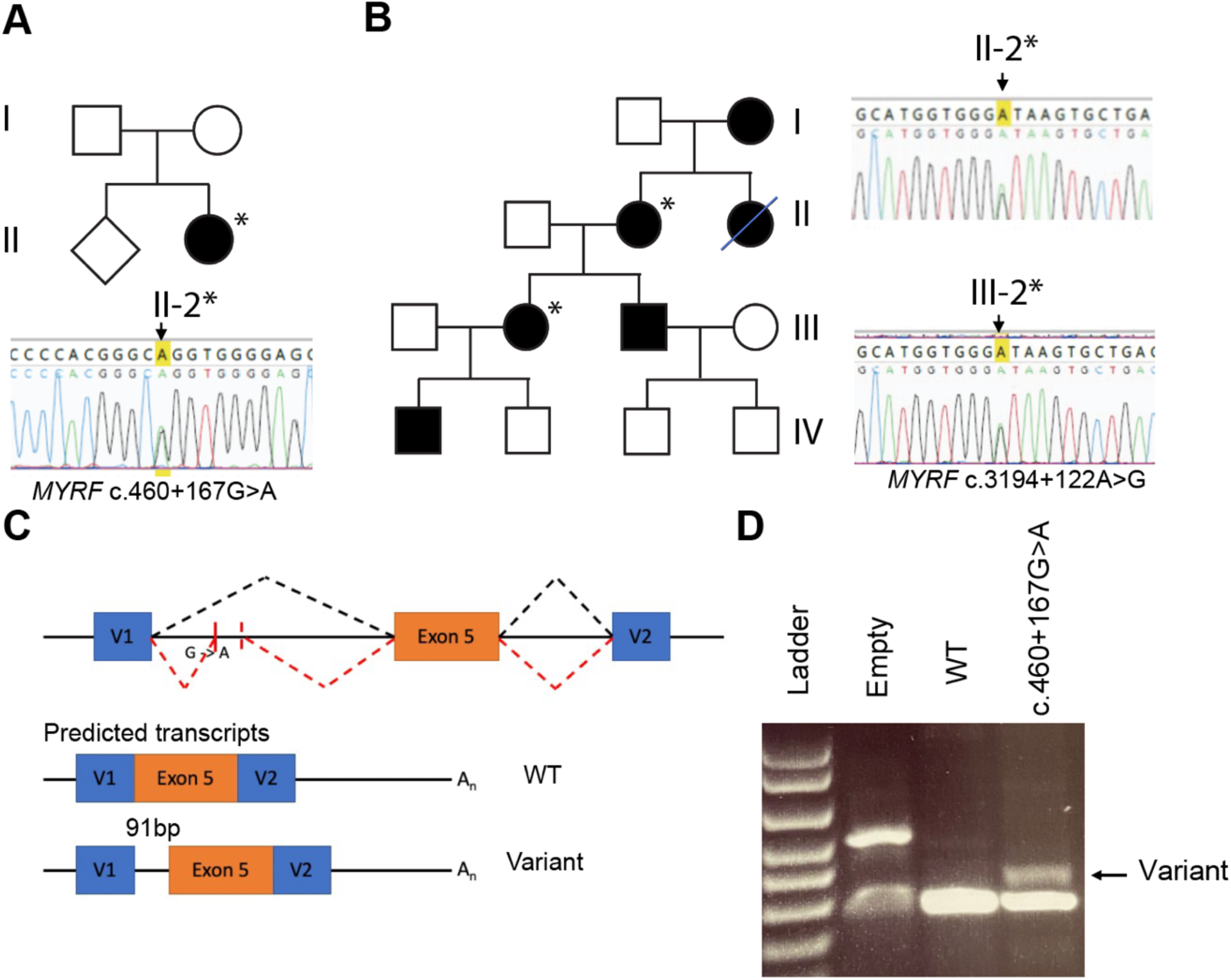
Novel deep intronic variant of *MYRF* alters splicing of mRNA transcript. (A-B) Two families were discovered with novel intronic variants in *MYRF*. (C) Schematic of the minigene RNA splicing assay and predicted products. (D) Minigene for variant c.460+167G>A shows altered splicing of MYRF at the mRNA level.

Clinical examination of family members revealed very short axial lengths in both families and classic features of nanophthalmos including choroidal folds and retinal detachment (Supplemental Figure 9, Supplemental Table 1). P01965 was a sporadic (Figure 6A), while family 2 exhibited a pattern of autosomal dominant inheritance, with demonstrated segregation in 2 generations. No systemic features of MYRF cardiac-urogenital syndrome were noted in any of the affected individuals or other family members.

The high spliceAI (33) scores for the variants in family 1 and 2 (c.460+167G>A, score = 0.992 and c.3194+122A>G, score = 0.8, respectively) indicate that the variants have a high probability of altering splicing (Supplemental Table 2). Predicted splicing affects for the first variant included gain of a cryptic acceptor site (c.460+167G>A), leading to a predicted pseudo-exon insertion of 91 bp (Figure 6C) and a frameshift leading to either early protein truncation or loss of transcript via nonsense mediated decay. An exon trap minigene splicing assay confirmed that the c.460+167G>A variant altered splicing compared to WT *MYRF* (Figure 6D). The variant preserved some normal splicing (283 bp WT exon 5 amplicon) along with a 374 bp fragment corresponding to the predicted pseudo-exon inclusion (Figure 6D). To quantify the fraction of transcripts that have aberrant splicing, PCR fragments were TA cloned, and individual colonies were sequenced (n=18). 7/18 (38.9%) contained RNA produced from the cryptic splice site usage, while the remaining used the normal splice acceptor. For Family 2, we confirmed co-segregation of the variant in two generations, yet we did not detect pseudo-exon inclusion using the exon trap vector system (Supplemental Figure 10). The variant may have a cell-type specific or contextual effect on splicing not detected by our assay or affect *MYRF* function through a different mechanism. WGS did not identify any other potential causes of nanophthalmos in this pedigree. Together, these data suggest that intronic splice variants in *MYRF* may be another mechanism leading to the pathogenesis of nanophthalmos and isolated ocular phenotypes may be driven by partial loss-of-function alleles.

## DISCUSSION

Though originally identified as master regulator of oligodendrocyte maturation, growing evidence supports a role for MYRF in regulating RPE development, differentiation, and maturation (11, 12). Our previous studies underline the importance of MYRF in the developing RPE showing that conditional loss of *Myrf* in mice can be disastrous for RPE and retinal development (11). Interestingly, there are a cluster of variants located in the conserved C2 domain that manifest as isolated ocular diseases (8). Our data support the idea that these variants are hypomorphic alleles, and other partial loss-of-function alleles, including splicing variants, may contribute to isolated ocular disease by altering gene dosage of *MYRF* (Figure 7).

**Figure 7.**
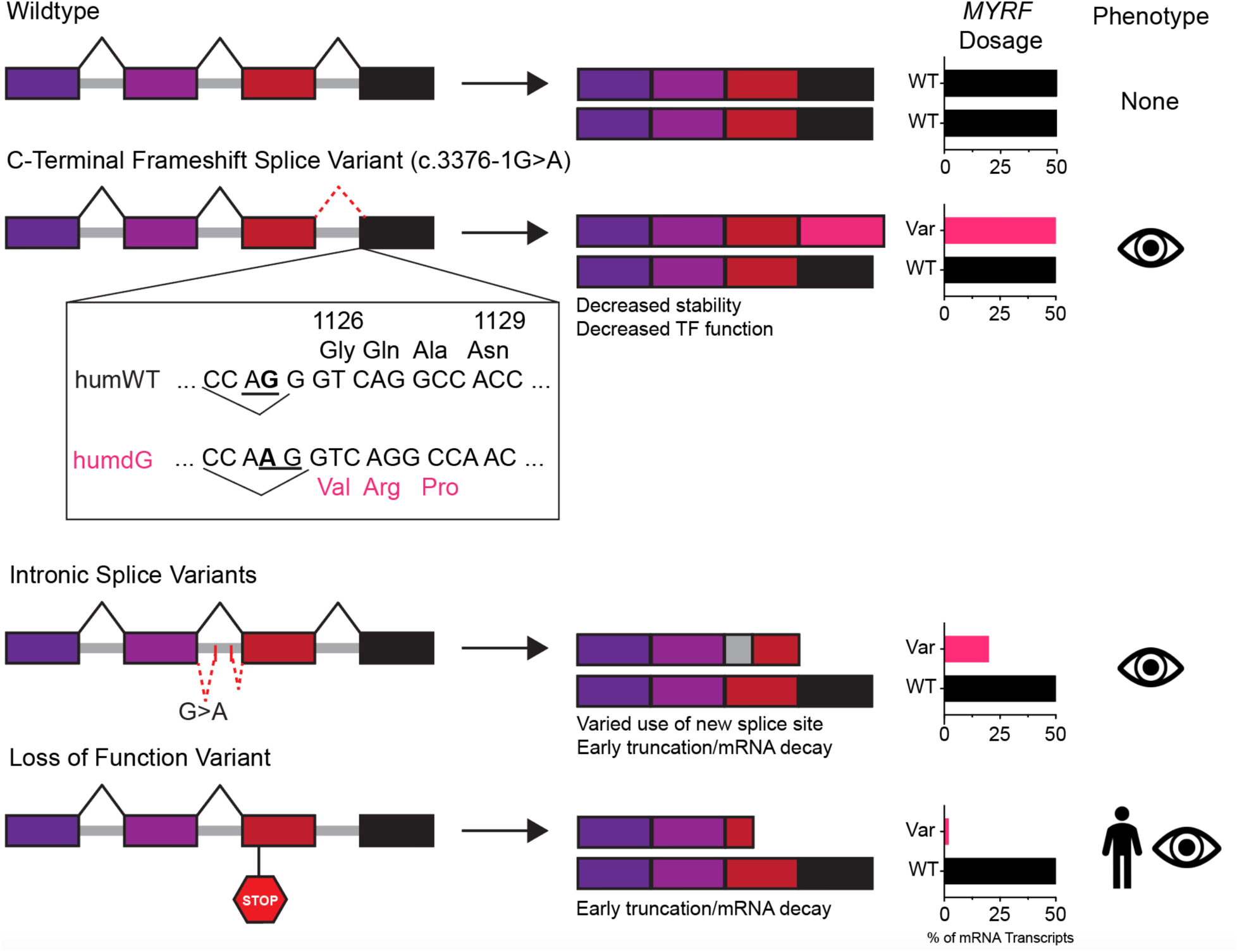
Working model of *MYRF* splice variant functional and phenotypic effects contributing to isolated ocular and syndromic disease pathogenesis. Wildtype *MYRF* retains transcription factor function resulting in no phenotypic effects. A C-terminal frameshift splice variant results in a 31 amino acid frameshift leading to decreased C-terminus stability and transcriptional activation, leading to reduced transcription factor function and isolated ocular disease. Intronic splicing variants lead to frameshift that results in early truncation or non-sense mediated RNA decay. These splice sites are only partially used, leading to modest decrease in MYRF transcription factor function and isolated ocular disease. In contrast, complete loss of function alleles either lead to early truncation/non-sense mediated decay or alter critical functional domains (i.e. DNA binding, cleavage) and lead to severe, syndromic symptoms.

### C-Terminal *MYRF* Variant Acts as a Loss of Function Allele

Several lines of evidence support that the C-terminal MYRF variant is a partial loss of function allele. First, embryonic lethality is observed in mice harboring a homozygous C-terminal truncating allele in *Myrf,* in line with the autosomal dominant inheritance pattern seen in the human disease. Embryonic lethality is observed later in gestation than constitutional knockout of *Myrf*, with skewed Mendelian ratios starting at E17.5, rather than lethality during cardiac development midgestation (34). Second, known transcriptional targets and similar pathways are dysregulated in our *Myrf^humdG/humdG^* mouse model, including ECM, TGFß signaling, Wnt signaling, structural regulation, and RPE maturation pathways, and *Myrf* itself. Third, transcriptional and phenotypic effects in *Myrf^humdG/humdG^* were milder than in comparable *Rx>Cre Myrf^fl/fl^* mice. *Myrf^humdG/humdG^* had normal appearing RPE pigmentation, modest reduction in *Tmem98* expression, and lower magnitude of DEGs among shared pathways with the *Rx>Cre Myrf^fl/fl^* eyes. Additionally, no phenotypic changes in the heterozygous *Myrf^humdG/+^* eyes over 1 year and the direct comparison of scRNA-seq datasets with the conditional knockout suggest that the C-terminal *MYRF* variant allele retains some partial function. Although molecular changes are picked up on scRNA-sequencing data, the phenotypic changes in the *Myrf^humdG/humdG^*are more modest in comparison to the *Rx>Cre Myrf^fl/fl^* mice. For example, genes in the pigmentation pathway are present in the downregulated DEGs of *Myrf^humdG/humdG^* such as *Oca2* at E17.5, but these mice show no signs of depigmentation at embryonic stages. Fourth, the C-terminal frameshift *MYRF* allele exhibits only subtle processing defects, with decreased steady state stability and transcriptional activity in vitro, but no differences in protein, localization, or cleavage in the context of an overexpression assay.

A previous study developed a mouse model to investigate a similar variant, *MYRF* c.3260delG/p.(Gly1087Valfs*151) mutation, which corresponds to the human *MYRF* c.3377delG/p.(Gly1126Valsf*31) mutation (13). This mutation causes the same C-terminal frameshift in humans as the c.3776-1G>A discovered by our lab. While there was support for loss of function for this allele, with lower mRNA and protein expression of MYRF(13), the mouse allele does not mimic the human allele well given the difference in the frameshift amino acid sequence and the 3’UTR sequence. Further, comparison of embryonic lethality of the Gly1087Valfs*151 variant allele to our current study is not possible, as the focus of the prior study was on only heterozygous mice. As such, our humanized mouse has more fidelity to model the human disease by replacing the last exon with human WT or human variant DNA sequence and recapitulates the 31 amino acid frameshift in patients. Some discrepancies remain between the models as we did not observe evidence to support changes in thickness of GCL or INL layers (measured by SD-OCT), phenotypic/morphological changes in heterozygous mice, or signs of primary angle closure glaucoma. These may be related to the different nature of the allele, CRISPR off-target effects, or strain/background specific effects. Our humanized mice and wildtype mice exhibit mRNA expression exclusively within the RPE cluster, therefore antibody detection of MYRF in other eye structures may represent cross-reactivity or background (11, 12). Our data, together with these previous models, support that the assertion that C-terminal *Myrf* variant is acting as a partial loss of function allele. However, it is important to note that future studies are necessary to completely rule out the possibility of the C-terminal variant acting via dominant negative interactions with the wildtype MYRF, as these mechanisms could not be assessed easily in our animal model or overexpression cell culture models.

### Consequences of Functional Loss of C-Terminal Domain

Our results show that C-terminal MYRF variant can cleave and localize properly to the nucleus like the WT protein. This is consistent with previous work that demonstrated a truncated version of MYRF lacking the C-terminal domain (MYRF-1:756) maintains proper homotrimerization and processing in oligodendrocytes (2). Previous work has also shown that homotrimerization is necessary for MYRF self-cleavage (1). This implies that the C-terminal variant is also able to form the homotrimer structure necessary to facilitate cleavage of MYRF. As the C-terminal MYRF variant homotrimerizes and cleaves, we might expect it to be capable activating transcription of its downstream targets. However, our scRNA-seq and qRT-PCR data show downregulation of key target genes and pathways involved in RPE development including the Wnt signaling, TGFß signaling, pigmentation, structural organization and epithelial cell development. The changes in these pathways are consistent with many changes detected in our conditional knock out model. For example, top downregulated hits in MYRF C-terminal variant RPE included known eye size disorder genes. Serpine3 encodes a serine protease inhibitor, which has been shown to be significantly upregulated in induced myopia models in zebrafish and chicks (28, 29). In contrast, it is significantly reduced our model, suggesting it may play a bidirectional role in eye size regulation like *Prss56*. Additionally, other top downregulated genes *Ankfn1* and *Kcnq5* are known GWAS hits for refractive error and/or myopia (35–37). Downregulated *Ankfn1* mRNA expression has recently been discovered as a defining feature of healthy vs. dedifferentiated RPE in mouse models exposed to cigarette smoke exposed or aged (27). *Kcnq5* encodes a voltage gated potassium channel expressed in the RPE (human, primate, and bovine) and believed to be involved in active transport of K^+^ from the retina to the choroid (30, 31). This data suggests that the C-terminal variant is likely influencing expression of genes involved in eye size regulation. Interestingly, although dG-MYRF maintains normal protein processing, at steady state the full length and C-terminal fragments are less stable. It is possible that the instability observed at steady state levels may be a driving factor behind decreased transcriptional activation of downstream target genes. Further, in the *in vitro* overexpression context, we see loss of *MYRF* autoregulation and inability to upregulate the TMEM98 downstream target the *dG-MYRF* mutant. These results are consistent with other studies supporting a role for the C-terminus of MYRF in regulation of its function, as removal of the ER-resident portion in *in vitro* assays results in decreased levels of target gene transcription (2, 38). Furthermore, overexpression of an mouse *Myrf* plasmid missing the ER portion results in decreased expression of genetic markers of differentiation in maturing oligodendrocytes *in vitro* (2). Together these data suggest that the C-terminal MYRF variant and removal of the ER portion may decrease target gene expression even though the protein appears to maintain normal cleavage and localization dynamics in cell culture, suggesting that other important protein-protein interactions may be at play.

### Role of C-Terminal Domain in ER Homeostasis

The function of the C-terminal domain that remains within the ER is not well understood. High grade pancreatic cancers have higher MYRF levels, altered ER structure, and improved ER homeostasis (38). Like cancer cells, RPE cells are highly metabolically active and rely on ER health for normal function (39). Results from scRNA-sequencing GO term analysis suggest that presence of the *Myrf^humdG^* causes increase gene expression in translation, ribosomal processes, and glucose metabolism pathways. This supports the speculation that dG-MYRF is no longer able to fulfill its role maintaining ER homeostasis and the RPE may be attempting to upregulate these processes as a compensatory mechanism. Together, the findings of discordant DEGs and pathways in *Myrf^humdG/humdG^* and *Rx>Cre Myrf^fl/fl^* eyes suggest potential for non-transcriptional roles of the MYRF C-terminus, though further studies are needed to fully explore this.

### Presence of Splice Altering Variants in Nanophthalmos

In addition to variants in well-characterized domains such as the DNA binding domain and the ICA cleavage domain, which fundamentally alter MYRF’s function to act as a transcription factor, we found predicted splice variants in *MYRF* that are associated with nanophthalmos. We discovered two novel, deep intronic variants in *MYRF* that are predicted to alter splicing, and one of these was confirmed by a minigene splicing assay. Our confirmed splicing variant of *MYRF* is expected to result in an early frameshift and produce a nonfunctional protein product. The minigene splicing assay suggests that pseudo-exon inclusion occurs in ~40% of the transcripts. These data together with our animal and cell culture evidence from the C-terminal frameshift variant, support a tissue-specific, dosage-sensitivity threshold, whereby partial loss of function variants may lead to ocular disease only, while complete loss-of-function or dominant negative alleles may lead to syndromic disease. Dosage sensitivity of MYRF is supported by previous studies showing intrafamilial phenotypic variability (11, 17, 40). Indeed, most individuals with cardiac-urogenital syndrome that are phenotyped for hyperopia do have ocular abnormalities (3). As whole genome sequencing is applied more often, deep intronic variants in *MYRF* and other nanophthalmos-associated genes may be found (41).

In this study, we have investigated mechanisms by which MYRF alleles may cause isolated ocular disease. Our results emphasize the role of the C2 domain in regulation of the activity of MYRF in the RPE, as its alteration leads to decreased steady state stability, transcriptome wide dysregulation, and downregulation of pathways important for RPE development. We also demonstrate that the C-terminus of MYRF is important and necessary to produce viable offspring in mice. These results build upon the limited knowledge of the function and importance of the conserved C2 C-terminal domain of MYRF and offer steppingstones to further understand how C-terminal variants of MYRF contribute to disease pathogenesis. Further, our identification of variant alleles that alter splicing but maintain some appropriate splicing supports the conclusion that the eye is particularly sensitive to reduced MYRF function.

## METHODS

### Sex as a biological variable

Our study examined male and female animals, and similar findings are reported for both sexes.

### Plasmid Generation and Lentiviral Transduction ARPE-19 Cells

DNA constructs were generated (Twist Bioscience) in pLentiLox-IRES-GFP backbone vector of human HA-tagged WT-MYRF. The C-terminal MYRF mutant and the V679A cleavage deficient were generated by site-directed mutagenesis of the pLentiLox-FLAG-MYRF-IRES-GFP plasmid using partially overlapping primers (Forward: 5’-GCACTGCTGGTCAGGCCAACTGCAGTTCAGAGG-3’, Reverse: 5’-TGGCCTGACCAGCAGTGCCACCCGAAAGTGGTA-3’) and the Quik Change II Kit (Agilent, #200517). Initial mutagenesis junctions were verified via Sanger sequencing, and whole plasmid sequencing was confirmed by Nanopore sequencing (Plasmidsaurus). These bicistronic vectors contained an N-terminal FLAG tag used to detect both cleaved N-MYRF and full length MYRF as well as an eGFP reporter to measure transduction efficiency. Lentiviral constructs were packaged by the University of Michigan Vector Core (10X concentrated virus) using standard methods with the plasmids above. Viral titers were assessed by flow cytometry for GFP on sub-confluent ARPE-19 cells using a range of viral supernatant dilutions (10^-1^ to 10^-4^) with assistance from the University of Michigan Flow Cytometry Core.

### Protein Localization Studies

ARPE-19 cells were seeded at 40,000 cells/well on Nunc Lab-Tek II 4-well chamber slides. After 24 hours, cells were transduced with 10^-2^ dilution of WT-MYRF, C-terminal mutant MYRF, or V679A (10X concentrated) vectors in the presence of 100mg/mL LentiBoost (Mayflower Bio, # SB-P-LV-101-11). Images of GFP signal were taken on an inverted microscope (Thermo Fisher, EVOS FLc) 72 hours after lentiviral transduction. Cells were fixed with 4% PFA in 1XPBS for 10 min. Fixed cells were incubated with anti-FLAG 1:1000 (CST, #14793) and anti-Calnexin 1:200 (Sigma, #C4731) primary antibodies overnight at 4C. Fixed cells were then incubated with anti-goat rabbit Alexa Fluor 488 1:500 (Jackson ImmunoResearch Labs, # 111-545-144) and goat anti-mouse Alexa Fluor 555 1:1000 (Invitrogen, A-21422) secondary antibodies for 2 hours at RT. Cells were washed 3 times (5 min each) with 1XPBS and stained with DAPI. Slides were mounted with gold anti-fade mounting media (Thermo Fisher, P36930). Stained cells were then imaged at 63X using Leica SP8 confocal microscope.

### Protein Stability & Cleavage Studies

ARPE-19 cells (ATCC, #CRL-2302) were seeded at 40,000 cells in 24-well tissue culture plates. After 24 hours, sub-confluent cells were transduced with lentiviral vectors carrying WT-MYRF, C-terminal mutant MYRF, or V679A (MOI 3 from 10X concentrated stocks) vectors in the presence of 1mg/mL LentiBoost (Mayflower Bio, # SB-P-LV-101-11). Images of GFP reporter were taken on an inverted microscope (EVOS) 72 hours after lentiviral transduction. After 72 hours, cells were treated with 300ug/mL cycloheximide (Sigma Aldrich, Catalog #: C4859-1ML). Cell lysates were collected at 0, 12, 16, 20, 24 hours. Cells were rinsed twice with PBS and lysed with Pierce RIPA Buffer (Thermo Scientific, #89900). Protein concentration of cell lysates was quantified using Pierce BCA Protein Assay Kit (Thermo, #2325). Samples were denatured using 4X Laemmli Sample Buffer (Bio-Rad) supplemented with 2-Mercaptoethanol and boiled at 95C for 5min. Upon SDS-page, samples were transferred onto Immobilon-FL PVDF membrane (Millipore Sigma) and probed chicken anti-GFP 1:5000 (Abcam, #13970) or rabbit anti-GAPDH 1:1000 (Millipore Sigma, #ABS16) and rabbit anti-FLAG 1:1000 (CST, #14793) or mouse anti-MYRF^393-766^ (Ben Emery lab) primary antibodies. IRDye 800CW goat anti-rabbit IgG 1:20,000 (LICOR, 926-32211) and IRDye 680RD donkey anti-chicken IgG 1:20,000 (LICOR, #925-68075) were used as secondary antibodies. Blots were imaged using the Odyssey CLx and quantified using ImageStudio.

### Quantitative RT-PCR

For ARPE-19 experiments, cells were seeded at 50,000 cells/well on 24-well plates. After 24 hours, cells were transduced with 10^-2^ dilution of WT-MYRF, C-terminal mutant MYRF, or V679A (10X concentrated) vectors in the presence of 100mg/mL LentiBoost (Mayflower Bio, # SB-P-LV-101-11). After 72 hours, RNA was isolated from cells using the RNAqueous™-Micro Total RNA Isolation Kit (Invitrogen) according to the manufacturer’s protocol. cDNA was generated using the SuperScript III system (Thermo Fisher Scientific) following the recommended protocol, with no reverse transcriptase (RT) negative controls. Quantitative RT-PCR was performed on the Applied Biosystems 7500 Real Time PCR System using the Taqman Assay system with inventoried probes for *MYRF*, *TMEM98, B-ACTIN* (Thermo Fisher Scientific). Critical cycle threshold levels of each sample were normalized to levels of beta-actin. Fold activity was calculated using the ddCt method and reported relative to non-transfected cells.

For animal experiments, eye tissues were harvested from E15.5, E18.5 mouse embryos or 8-month-old mice. The lens, cornea, and optic nerve were dissected, and RNA was isolated using the RNAqueous™-Micro Total RNA Isolation Kit (Invitrogen) according to the manufacturer’s protocol. cDNA was generated using the SuperScript III system (Thermo Fisher Scientific) following the recommended protocol, with no reverse transcriptase (RT) negative controls. Quantitative RT-PCR was performed on the Applied Biosystems 7500 Real Time PCR System using the Taqman Assay system with inventoried probes for Myrf and Tmem98 (Thermo Fisher Scientific). Critical cycle threshold levels of each sample were normalized to levels of Hprt. Fold activity was calculated using the ddCt method and reported relative to control littermates.

### In Silico Modeling

PONDR (23) and AlphaFold2 (22) were used to in silico model changes in protein stability and structure WT-MYRF and dG-MYRF conserved C2 domain amino acid sequence. PONDR was used to predict differences in intrinsically ordered and disordered regions. VXLT combines three previous neural network models (VL1, XN, and XC) trained on NMR and x-ray crystallography data with varying lengths of disordered regions. VSL2 combines two predictors optimized for long (>30 residues) and short (<=30) disordered regions (20, 21, 23, 42). Alphafold2 was used to de novo predict 3D structure of the C2 domain in the presence or absence of the C-terminal frameshift variant amino acid sequence.

### Generation of Humanized *Myrf* alleles in Mice

Mice were genetically engineered using CRISPR-Cas9 replacing mouse *Myrf* exon 26 with the homologous human *MYRF* exon 27 and fused with mouse exon 25, including the C-terminal coding region the human 3’UTR sequence (Supplemental Figure #). This humanized *Myrf* allele (*Myrf^humWT^*) was generated by injecting fertilized eggs from C57BL/6J matings with a pair of spCas9 guide RNAs (25ng/µl each) (gRNA, PAM=NGG) that are 253 bps apart in outward orientation together with the spCas9 protein (NEB, 30 ng/µl). Sequences of the gRNAs are listed below: the upstream gRNA 5’GCTCACCAGCAATGTCACC3’ and the downstream gRNA 5’GGCAGTGCCACAGCC GGGAC3’. A single stranded oligo of 327 bps (IDT Megamer) carrying the human Exon27 sequence and homology arms of 100 bp each is also included in the injection mixture as the repair template for single strand DNA mediated homologous recombination (HR). The sequence of this HR template is 5’TAGGACAAGGCTGCCCTGACCTCTGTCTTCCCTGTCTCTCTAGGGCACCTCTCATCAGTGGCCAGTAACCATCCTGTCCTTCCGTGAATTCACATACCACTTtCGGGTGACATTGCTGgGTCAGGCCAACTGCAGTTCAGAGGCTCTCGCCCAGCCAGCCACAGACTACCACTTCCACTTCTACCGCCTGTGTGACTGAGCTGCCCTCCTGACAGTGCCACAGCCGcGACTGGAGTCCCTGGGCCCTCAACACTGGATGCAAATGTGTTACACTGGAGCCTGCTGCAGGCCAGCTCTCTGCTCTTCACTGCTTCCCTTGACTGGGGA3’ (327 bps). Founder mice were genotyped using primers (Forward: 5’-GTGGCTACCCTTGCTCTCAA-3’, Reverse: 5’-GGCGAGAGCCTCTGAAC TG-3’). We generated matched *Myrf^humdG^* mouse that recapitulates the splice site mutation found our previously described large familial cohort with nanophthalmos starting with heterozygous Myrf^humWT/+^, and using the following guide RNA (5’-CACTTCCGGGTGACATTGCT-3’) and ssDNA repair template: 5’-TGGCCAGTAACCATCCTGTCCTTCCGTGAATTCACATACCACTTCCGGGTGACATTGCTGGTCAGGCCAACTGCAGTTCAGAGGCTCTCGCCCAGCCAGCCACAGACTACCACTTCCACT-3’ *Myrf^humWT/humdG^*mice are viable and produce offspring, while *Myrf^humdG/humdG^* mice are embryonic lethal. Rx-Cre;*Myrf^fl/fl^* mice have been previously described (11). Mice were genotyped using standard methods (43) and primers listed above, followed by restriction digest with the BseYI restriction enzyme site to distinguish between the *Myrf^humdG^* and *Myrf^humWT^*alleles. Animals were also initially genotyped for *rd1*, *rd8*, and *rd10* according to established methods, to eliminate retinal degeneration alleles common in C57BL/6J mouse strains (44).

### RNAScope In Situ Hybridization

Heads were harvested from E15.5 embryos. Samples were fixed in 4% buffered paraformaldehyde (0.1 M NaPO_4_, pH 7.3) overnight at 4°C, dehydrated through increasing concentrations of ethanol up to 70%. The dehydrated samples were embedded in paraffin using the TissueTek VIP Model VIP5A-B1 (Sakura Finetek USA, Inc.) and the Shandon Histocentre 2 Model #64000012 embedding station (Thermo Fisher Scientific) and sectioned at 5-6μm. For *in situ* hybridization, RNAscope was performed using the RNAscope® Multiplex Fluorescent Detect V2 system (Advanced Cell Diagnostics [ACD], #323110). Briefly, paraffin was removed with two changes of xylene and then washed in 100% ethanol. Sections were treated with the hydrogen peroxide reagent for 10 minutes followed by two washes in distilled water. Target retrieval was performed by boiling in 1 X Target Retrieval Reagent for 7 minutes, followed by washing in distilled water and 100% ethanol. After drying the slides, the sections were treated with Protease Plus in prewarmed humidity chamber for 25 minutes at 40°C then washed in distilled water. Prewarmed RNAscope probes were applied, and slides were incubated in a humidity chamber at 40°C for 2 hours. The probes used were *Mm-Myrf* (ACD, 524061) and a negative control probe (ACD, 320871). After hybridization with the probe, sections were washed in 1 X Wash Buffer then incubated at 40°C with Amp1 reagent for 30 minutes, Amp2 reagent for 30 minutes, and Amp3 reagent for 15 minutes, with washes in 1 X Wash Buffer between each step. For signal development, the sections were then incubated in HRP-C1 for 15 minutes at 40°C, washed in 1 X Wash Buffer and then incubated with Cyanine 3, Opal-Tm 570 (Akoya Biosciences, FP1488001KT) fluorophore diluted 1:1500 in TSA plus buffer (ACD, 322809) for 30 minutes at 40°C. Sections were then washed in 1 X Wash Buffer then treated with HRP Blocker for 15 minutes at 40°C, stained with DAPI (Sigma, MBD0015) and mounted in ProLong Gold Antifade (Invitrogen, P36930).

### Retinal layer measurements, whole eye measurements, and cell counting

Eyes from 12-month animals were enucleated and aligned such that the optic nerve was in line with the central cornea such that both could be readily observed, and images were captured on the Leica MX10F dissecting microscope (Leica, Wetzlar, Germany). Ocular axial length was measured at 12-months using ImageJ 1.51m9 software (45), by marking a line from the central cornea to the base of the optic nerve. For the analysis, the following number of animals/eyes were used: *Myrf^humWT/+^* control, n=4 animals, 8 eyes; *Myrf^humdG/+^*, n=5 animals, 9 eyes.

### Mouse ocular imaging and electrophysiology

Eyes from 3-, 6-, 9-, and 12-month old control (*Myrf^humWT/+^*; n=8 eyes, 4 mice) and heterozygous mutant mice (*Myrf^humdG/+^*; n= 9 eyes, 10 mice) were evaluated sequentially in vivo by fundus photography, spectral domain OCT, and electroretinography as previously described (11). Data from *Myrf^humdG/+^*and controls were evaluated for statistical significance with one-way ANOVA in Graphpad Prism 10 (San Diego, CA, USA). If significant, subsequent pairwise comparisons were done with two-tailed Student’s t-test. A-wave and B-wave amplitudes from *Myrf^humdG/+^*eyes and control littermates were evaluated using two-tailed Student’s t-test in Graphpad Prism 10 (San Diego, CA, USA).

### RPE flat mounts

Eyes were enucleated from 12-month mice, and cornea, lens, optic nerve, and retina tissues were removed. At least 3 animals/eyes per genotype/age were used. Optic cups were fixed for 45 minutes in 4% PFA in 1XPBS. RPE samples were washed in PBS and blocked in 10% Normal Goat Serum (NGS), 1% BSA in PBTx for 2 hours. Primary antibody incubation occurred overnight at 4 °C with the antibodies and dilutions to the following antigens: TMEM98 (1:500, 14731-1-AP, Proteintech), Phalloidin (1:400, #A12380, Thermo Fisher). RPE samples were washed in PBS and incubated with Alexa-conjugated fluorescent secondary antibodies for 2 hours at room temperature. RPE samples were rinsed in PBS and the nuclei were stained with DAPI. A series of 3–4 radial cuts were made to flatten the RPE and underlying scleral and samples were mounted with Prolong Gold Antifade. Images were taken on a Lecia DM6000 compound microscope.

### REShAPE RPE Morphometric Analysis

RPE flat mounts (see above) were prepared from 12-month-old control (*Myrf^humWT/humWT^*; n= 4 eyes, 4 mice), heterozygous (*Myrf^humWT/humdG^*; n= 5 eyes, 10 mice) from a single cohort. RPE flat mounts were stained as above with Phalloidin to outline RPE cell borders. Whole RPE flat mounts were imaged using the LASX Navigator tiling software on a Lecia DM6000 compound microscope at 20X. Tiled images were fed into the REShAPE software (25) and processed producing a segmented image for each tile based on the phalloidin staining. Segmented images were further analyzed by the REShAPE software looking at morphometrics such as cell size, aspect ratio, hexagonality, and number of neighboring cells. Data points for RPE morphometrics from each tile were combined per biological replicate. Control and heterozygous RPE morphometrics were analyzed by performing a student’s t-test on the median value (cell size, aspect ratio, hexagonality, and number of neighboring cells).

### scRNA-seq Analysis

Eyes from E17.5 *Myrf^humWT/humWT^* (n=3 mice, 6 eyes) and *Myrf^humdG/humdG^* (n=3 mice, 6 eyes) were dissected removing the lens and optic nerve and leaving only the optic cup (retina/RPE/sclera/cornea) and dissociated using a solution containing 5 mM L-Cysteine (Sigma, C7352), 1mM EDTA (Sigma, E4884), 0.6 mM 2-mercaptoethanol (Sigma, M6250), and 1 mg/ml Papain (Roche, 10108014001). Eye cups were incubated at 37°C for 10 minutes with trituration at 2 minute intervals. Dissociation was stopped using Neurobasal Media (Invitrogen, 12348017) and 10% Fetal Bovine Serum (FBS) (Corning, MT35010CV), and cells were collected by centrifugation at 300 RCF for 5 minutes at 4°C. Single cells were resuspended in Neurobasal Media with 3% FBS. Eyes were processed for single cell RNA-sequencing (scRNA-seq) at the University of Michigan Advanced Genomics Core using manufacturer’s protocols for 10x Single Cell Expression 3’ library kit and sequencing on the NovaSeq (S4) 300 cycle (Illumina, San Diego, CA). Sequencing data was aligned, and count matrices and quality control results were generated using the CellRanger 7.1.0 pipeline. SEURAT 3.1.2 package (46, 47) was used to performed integration, unsupervised clustering, and cell type identification on scRNA-seq of E17.5 optic cups from this study with previously published scRNA-seq datasets from E15.5 and P0 RxCre;Myrf^fl/fl^ mouse optic cups and paired controls (12). Quality control thresholds were set to nFeature_RNA > 200 and percent.mt < 15. SEURAT 5.1.0 package was used to generate up and down regulated DEGs in the RPE cluster. The ClusterProfiler 4.14.6 (48) package were used to look for enriched GO terms and Gene Set Enrichment Analysis (GSEA) analysis (49). UCell was used to assign “scores” to enriched GO term and GSEA pathways in *Myrf^humWT/humWT^* (n=3 mice, 6 eyes) and *Myrf^humdG/humdG^*mice on a per cell basis (50).

### Patient Collection, Sequencing, and Analysis

The patient cohort consisting of individuals with axial length <21 mm or high hyperopia > +5.50 spherical equivalent was previously described (32). Clinical and biometric data were reviewed for patients. DNA from whole blood or saliva was extracted according to standard procedures as before. Patient were sequenced at the National Intramural Sequencing Center by Illumina short-read whole genome sequencing using the Illumina NovaSeq platform. Reads were aligned and variants were called and annotated via a customized pipeline (https://github.com/Bin-Guan/NGS_genotype_calling and https://github.com/Bin-Guan/variant_prioritization).

### Minigene splicing assays and patient blood RNA analysis

Two MYRF splicing variants were identified in families with nanophthalmos. A cryptic splice donor site (c.3194+12A>G) was identified in intron 24 of sample G04825 and a cryptic splice acceptor site (c.460+167G>A) was identified in intron 4 of sample G01965. G04825 and G01965 wild type and variant sequences were cloned into the pSLP3 exon trap vector. HEK293T cells were seeded in 6 well plates at a density of 75,000 cells per well and grown overnight. 970ng and plasmid DNA and 30ng GFP plasmid were transfected into HEK293T cells using Fugene 6 at a 3:1 ratio. Cells were incubated overnight and then harvested the next day for RNA isolation using the QIAGEN Mini RNA Isolation Kit. cDNA was generated using SuperscriptIII and random primers following manufacturer’s protocols. Splicing within the exon trap vector was analyzed by amplifying across the exon trap with for V1 primer 5’ TCTGAGTCACCTGGACAACC3’ and rev V2 primer 5’ ATCTCAGTGGTATTTGTGAGC 3’. PCR products were confirmed by Sanger sequencing. The PCR products were cloned into the TOPO TA cloning vector. 18 colonies were picked and sequenced to test the frequency of splicing into the cryptic splice acceptor site.

### Statistics

A chi-square test was used to detect significant skewing of expected Mendelian ratios of genotypes at E12.5-P0. A Wilcoxon Rank Sum test was used to identify DEGs between two groups of cells in the scRNA-seq dataset. For differential gene expression analysis, an adjusted p-value was calculated based on Bonferroni correction using all genes in the dataset. DEGs were considered significant with a p-adjusted value less than 0.1. ANOVA was used when comparing the means of three or more groups. A student’s T-test was used when comparing the means of only two groups.

### Study Approval

Human subjects work was approved by the Institutional Review Board (IRB) at the University of Michigan under protocol HUM00046159. All subjects provided written informed consent for their inclusion. Mouse studies were approved by the Institutional Animal Care and Use Committee at the University of Michigan under protocol PRO00010115 and PRO00011748.

## Data Availability

Single cell RNA sequencing data will be deposited in GEO (Accession number: ******). Primary data is provided for all the experiments. All additional relevant data can be found within the article and its supplementary information and primary data are available upon request from the corresponding author.

## Acknowledgements

We would like to acknowledge our funding sources: NEI K08 EY032098, the E. Matilda Ziegler Foundation for the Blind, Bright Focus Foundation (M2022011N), the Knight Templar Eye Foundation, Vision Research Core (NEI P30-EY007003), Shared Instrument Grant S10 (NIH SIG grant S10OD28612), NEI T32-EY013934, and NEI intramural funding. We would like to thank the Thom Saunders, Elizabeth Hughes, and the University of Michigan Transgenic Core for helping to generate our humanized mouse model. We would also like to acknowledge the Besirli lab (Eric Weh) for the modified REShAPE protocol and help with running tiles in REShAPE software. The authors are grateful to the Advanced Genomics Core for assistance with single cell RNA sequencing; to Ben Emery for helpful advice; to Sally Camper, Andrew Lieberman, Tom Wilson for helpful discussions.

