## Supplemental Materials for "Splicing variants in MYRF cause partial loss of function in the retinal pigment epithelium"

### SUPPLEMENTAL FIGURES

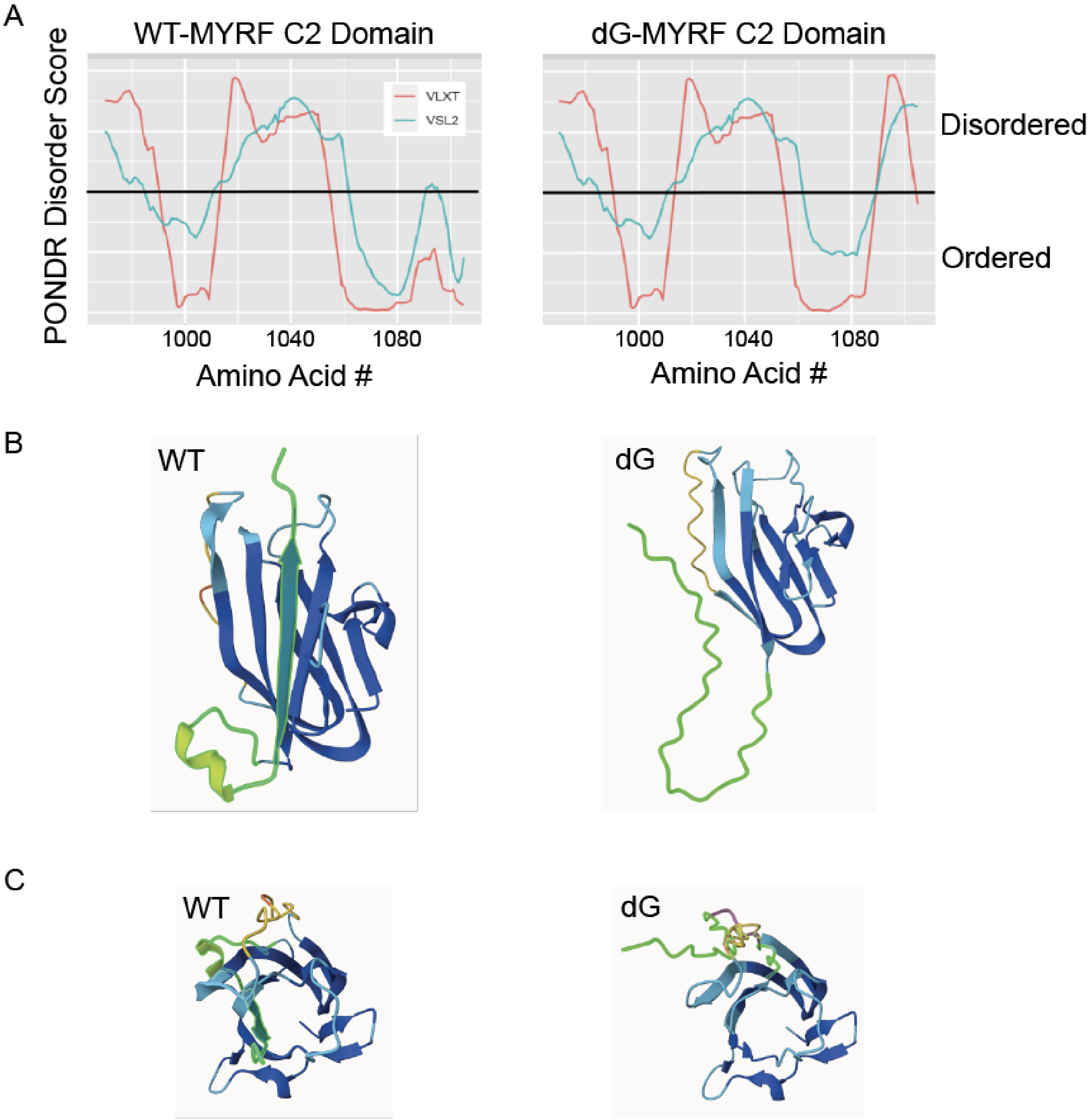


**Supplemental Figure 1. In Silico Modeling of MYRF C-terminal Variant Protein. (A)** The Predictor of Natural Disordered Regions (PONDR) algorithm suggests that the *dG-MYRF* variant creates a more disordered C2 domain than in *WT-MYRF*, PONDR score > 0.5 vs < 0.5, respectively. (**B-C**) AlphaFold2 de novo folding of the C-terminal domain of *MYRF* showing loss of beta sheet and alpha-helix structures as shown from the side (B) and from overhead (C). Region highlighted in green is the amino acid sequence affected by the C-terminal frameshift variant.

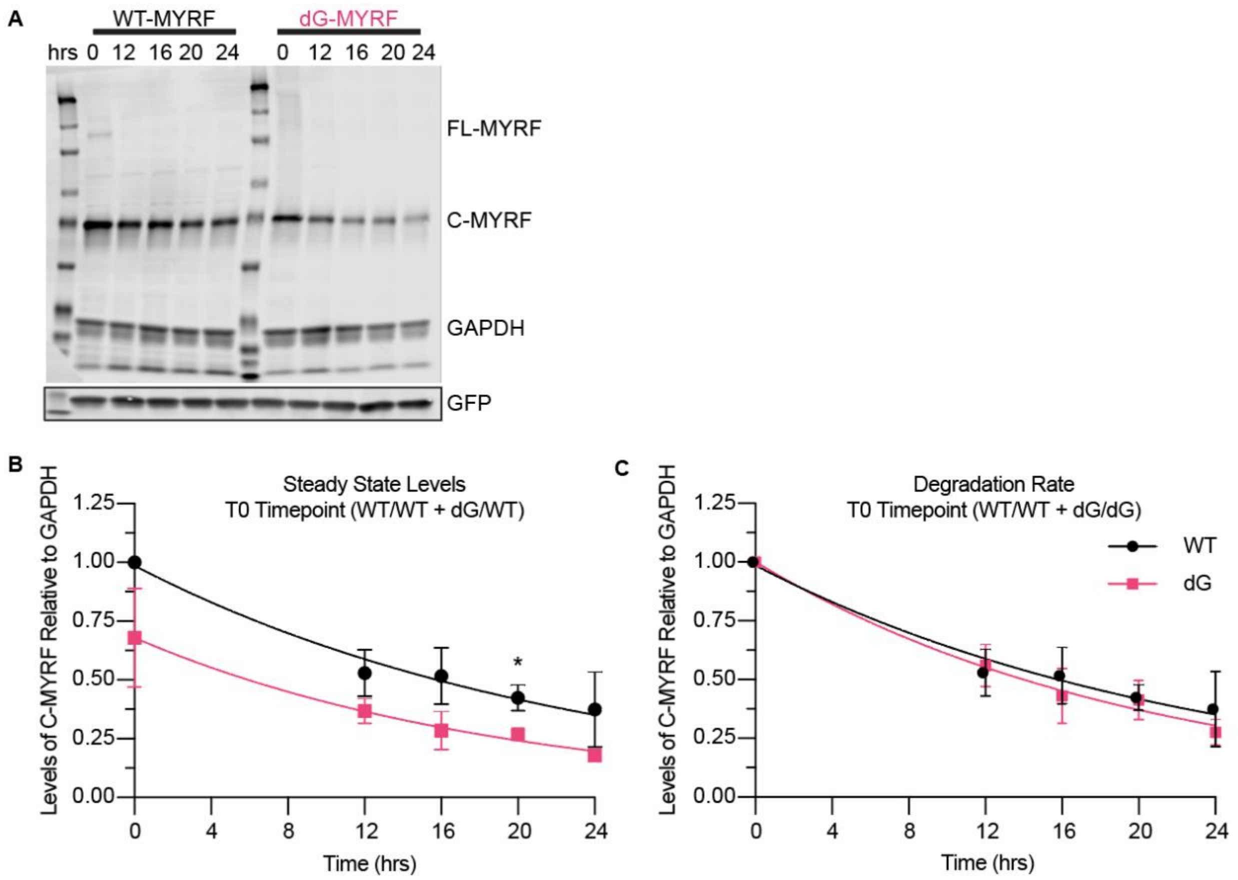


**Supplemental Figure 2. Cycloheximide pulse-chase assay detects reduced steady state levels of variant dG-MYRF.** (**A-B**) ARPE-19 cells were transduced with WT-MYRF and dG-MYRF. After 72 hours, cells were treated with 300ug/mL cycloheximide (CHX) to inhibit translation. Cell lysates were collected at 0, 12, 16, 20, 24 hours. Steady state levels (0 hr timepoint) of the of the full length (~140kDa) or C-terminal fragment (~70kDa) of MYRF in the dG-MYRF protein were decreased compared to WT-MYRF by Western blotting (A) and quantitated relative to GAPDH (*p<0.05) (B). (**C**) After adjusting the normalized values at time = 0 to 1, revealed no difference in the degradation rates of WT or dG-MYRF full length (~140kDa) or C-terminal fragments (~70kDa) over a 24-hour chase period (n=3).

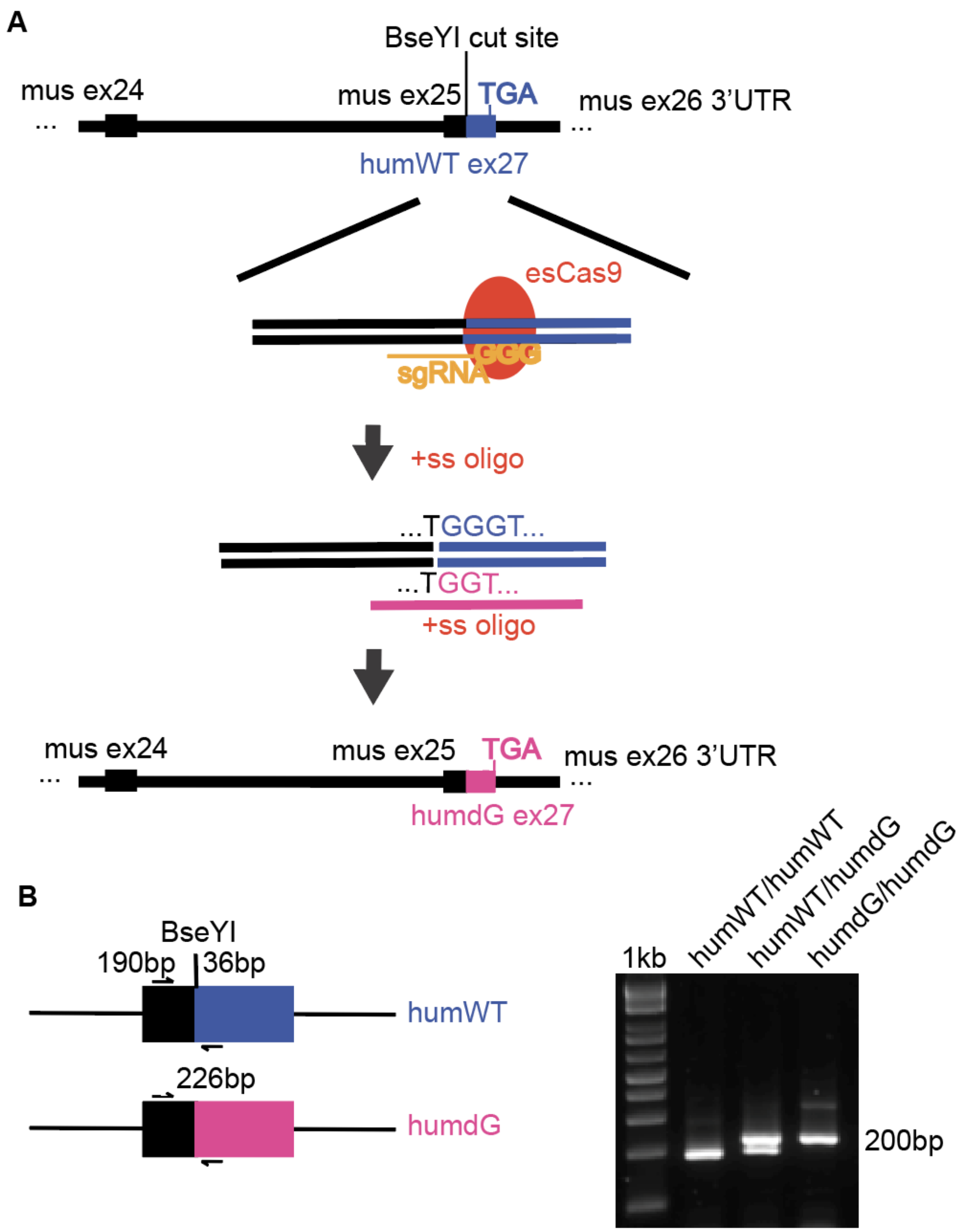


**Supplement Figure 3. Generation of the humanized C-terminal *Myrf* mouse model. (A)** CRISPR/Cas9 strategy to generate the humanized WT (humWT) and the C-terminal variant *Myrf* (humdG) alleles. (**B**) Genotyping strategy to differentiate humWT vs. humdG alleles using the BseY1 cut site which is lost in the humdG sequence.

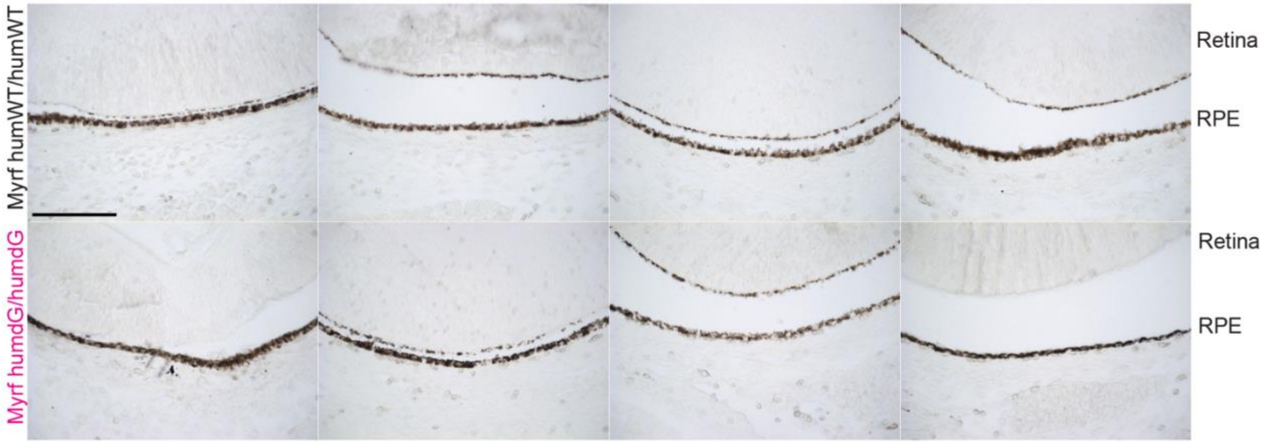


**Supplemental Figure 4. *Myrf*$^{humdG/humdG}$ show no signs of depigmentation at embryonic stages.** Brightfield images from E15.5-E16.5 embryos showing no change in pigmentation in the RPE layer of homozygous *Myrf*$^{humdG}$ embryos relative to *Myrf*$^{humWT}$ (scale bar = 100μm) (n=6-8 embryos per genotype).

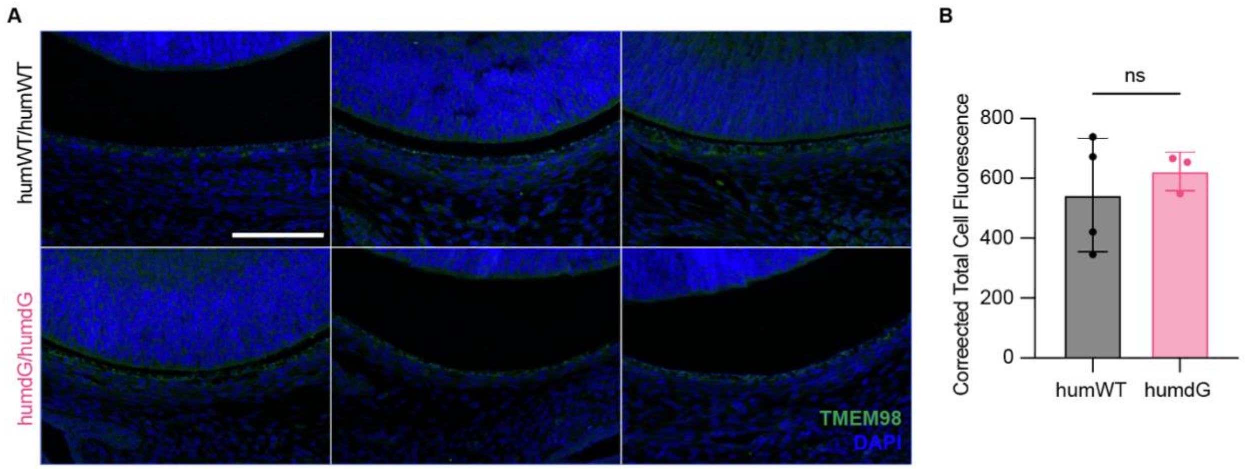


**Supplemental Figure 5.** ***Myrf*$^{humdG/humdG}$ show no reduction in TMEM98 protein at embryonic stages. (A)** IHC images of TMEM98 staining in E15.5 embryos showing no change immunoreactivity in the RPE layer of homozygous *Myrf*$^{humdG}$ embryos (scale bar = 100µm) (n=3-4 embryos per genotype). (**B**) Quantification of TMEM98 protein staining by corrected total cell fluorescence.

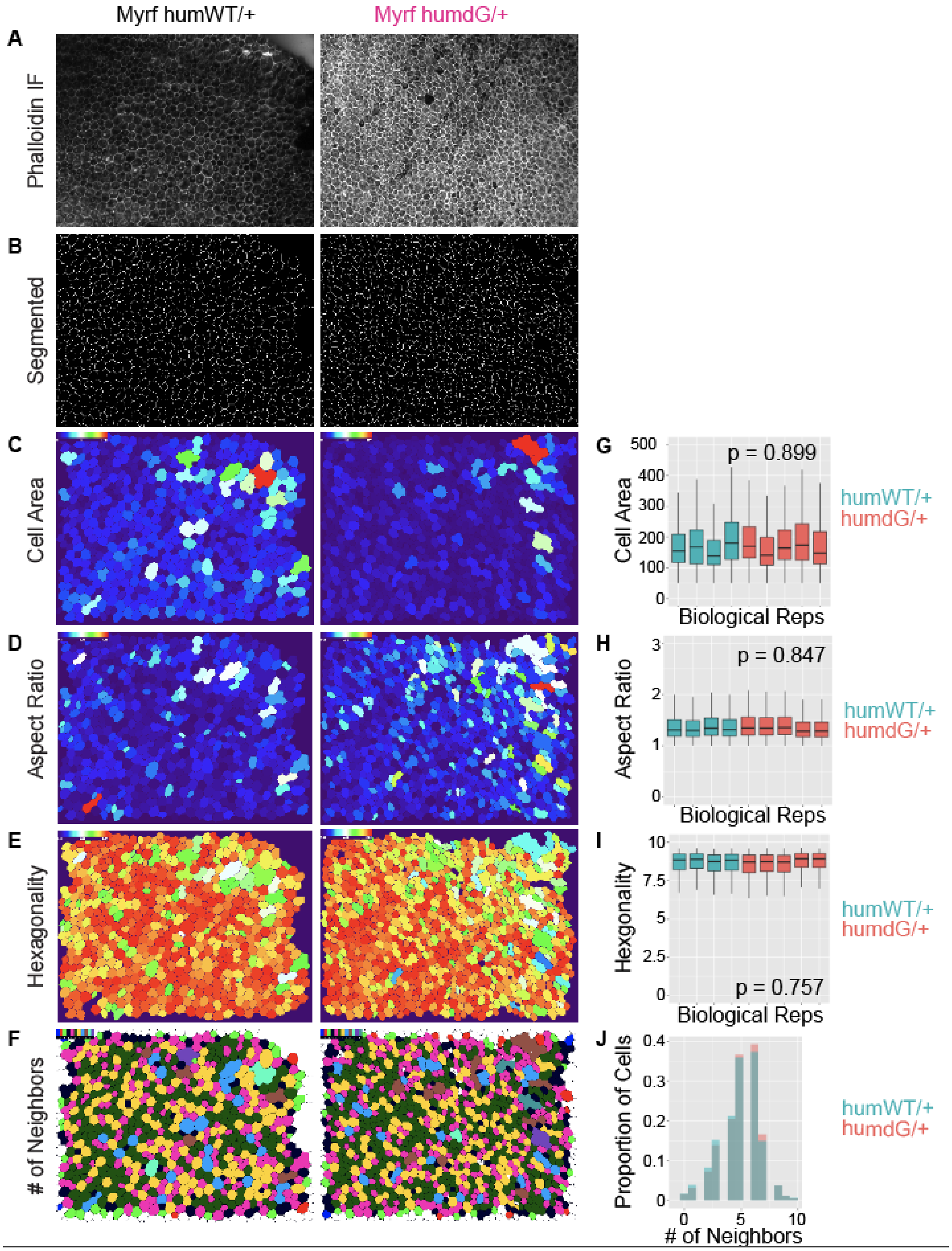


**Supplemental Figure 6. Morphometric analysis of *Myrf*$^{humdG/+}$ RPE reveals no differences in morphometrics.** (A-F) Representative output images from modified REShAPE AI software analysis examining segmentation (B), cell area (C), aspect ratio (D), hexagonality (E), # of neighbors (F) starting from an RPE flatmount stained with rabbit anti-phalloidin (1:400) to outline cell borders (n=4-5 per genotype). (G-J) Student's t-test showed no significant differences in compare median values for cell size (G), aspect ratio (AR, H), and hexagonality (I), and number of neighbors between genotypes.

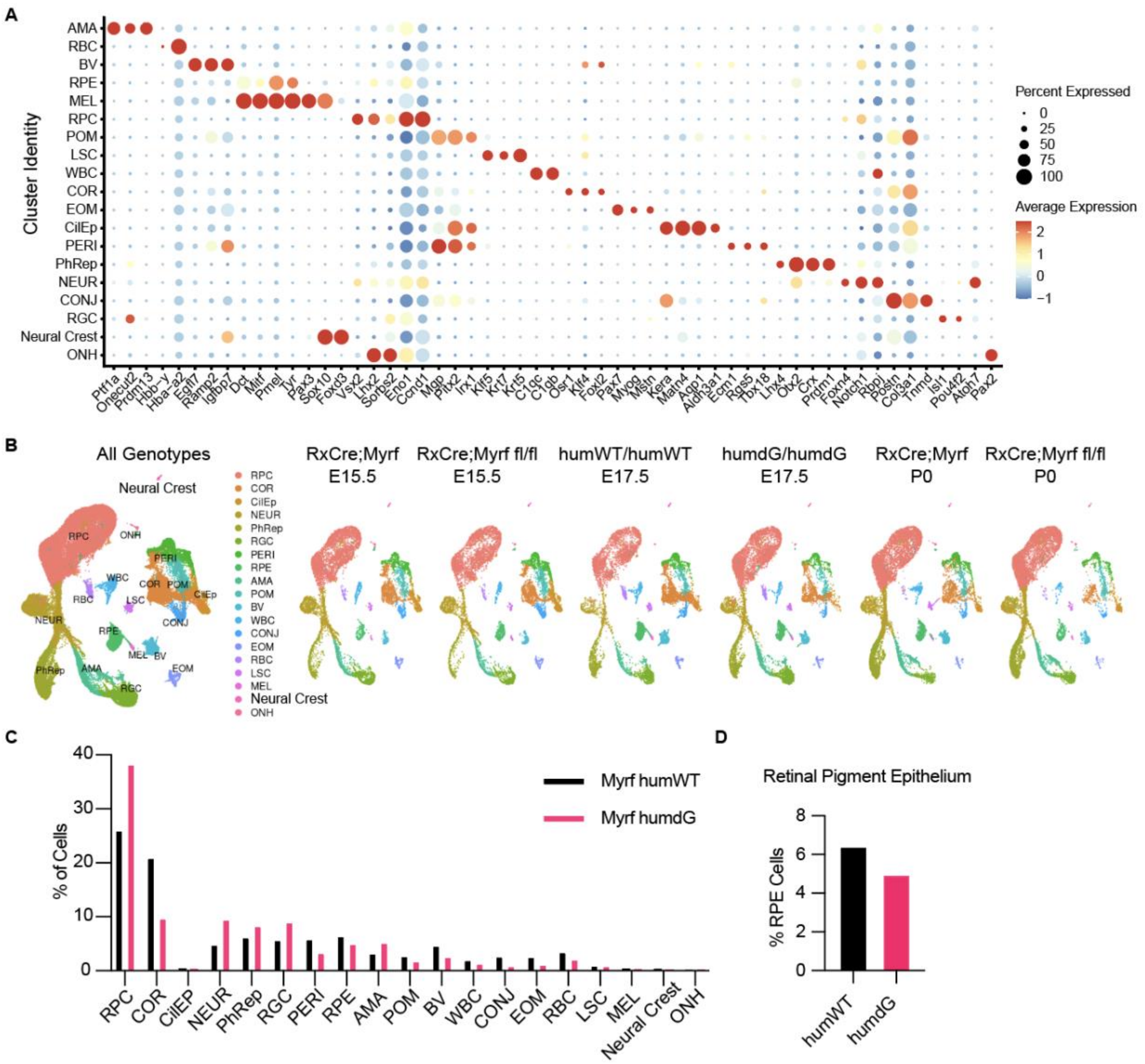

**Supplemental Figure 7. Identification of Optic Cup Clusters in Single Cell RNA-sequencing Dataset. (A)** Markers previously established in the field (12) were used for the identification of all major cell types within the optic cup. (**B**) Previously published *RxCre;Myrf*$^{fl/fl}$ E15 and P0 datasets (12) were integrated with the humanized *Myrf* datasets to improve clustering and used for cross comparison of downstream analyses. All datasets contained all major cell types, and no novel clusters were observed. (**C-D**) Percent of cells in each cluster by genotype (C). Percent of RPE cells in humWT (5%) compared to humdG optic cup (6.4%) (D).

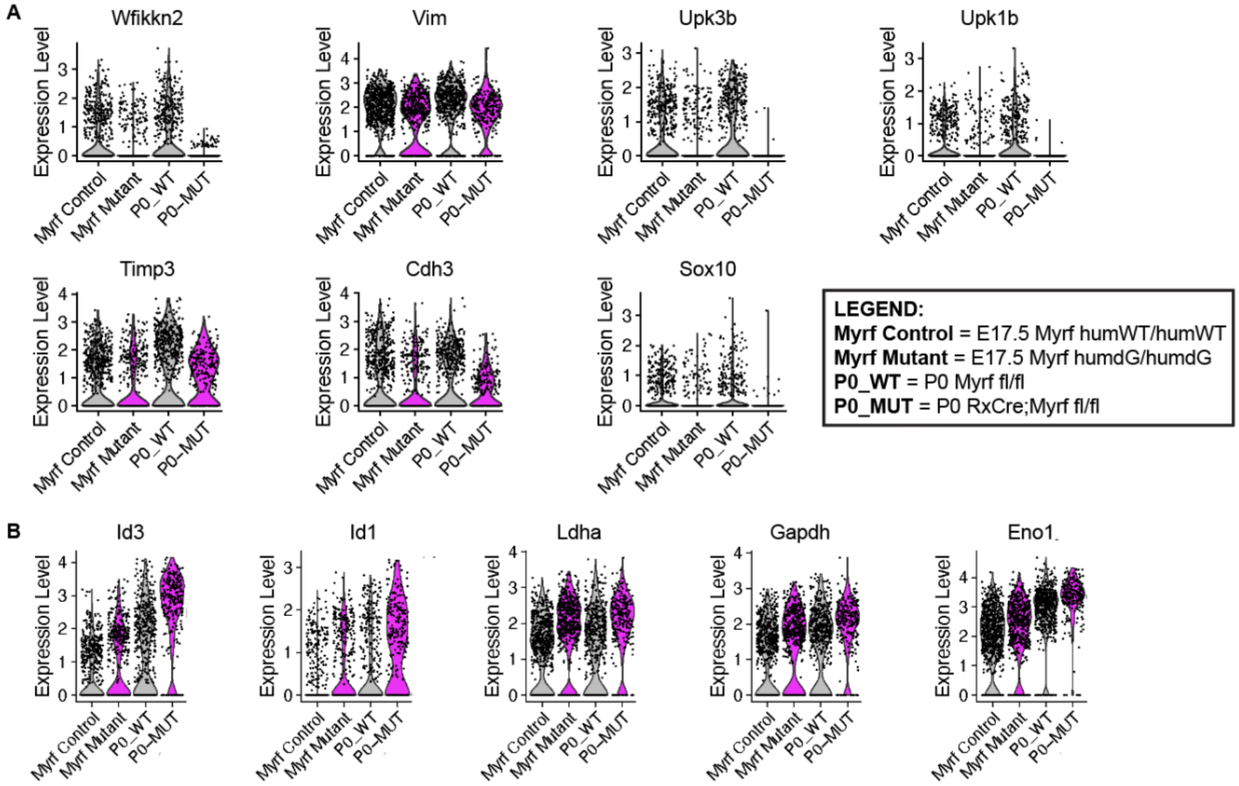


**Supplemental Figure 8. Concordance of Shared DEGs in Humanized C-Terminal Variant and Conditional Knock Out *Myrf* Mouse Models**. Downstream analysis showing violin plots with single cells plotted for selected differentially expressed genes in both *Myrf*$^{humdG/humdG}$ and *RxCre;Myrf*$^{fl/fl}$ mice was performed. Shared downregulated (A) and upregulated (B) genes related to RPE development and maintenance were all similarly altered in *Myrf*$^{humdG/humdG}$ and *RxCre;Myrf*$^{fl/fl}$ mice relative to their controls, but the magnitude of the effect was greater for some genes in *RxCre;Myrf*$^{fl/fl}$ relative to *Myrf*$^{humdG/humdG}$. For example, note *Id3* and *Id1.*

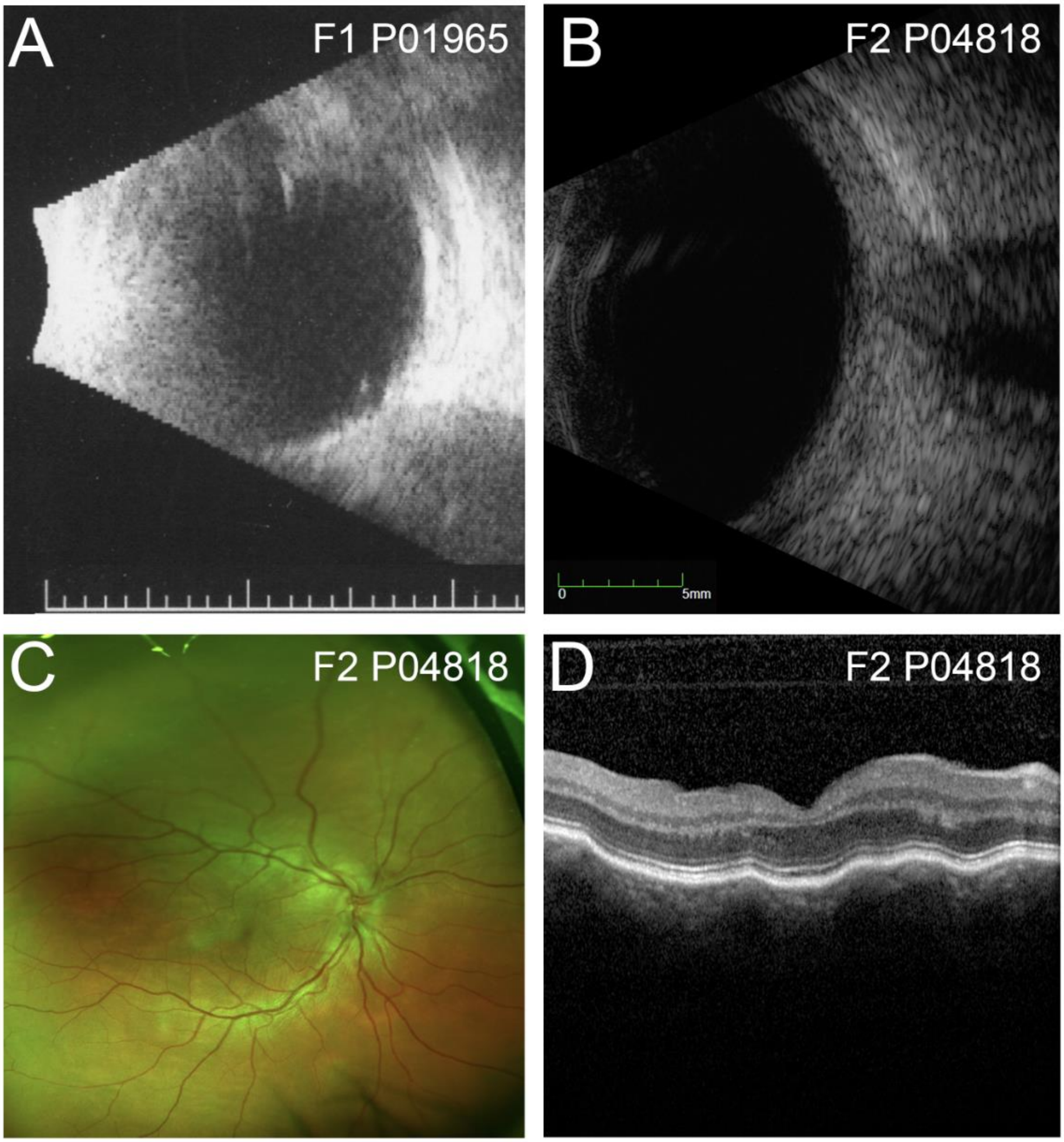


**Supplemental Figure 9. Clinical features of families carrying MYRF intronic variants.** (A-B) Ultrasound of right eyes of P01965 (A) and P04818 showing increased scleral thickness and reduced axial length 17.4 mm and 16.9 mm, respectively. (C) Optos photo of P04818 right eye showing vascular tortuousity, optic disc crowding, and no signs of retinal degeneration. (D) SD-OCT of P04818 showing choroidal folds.

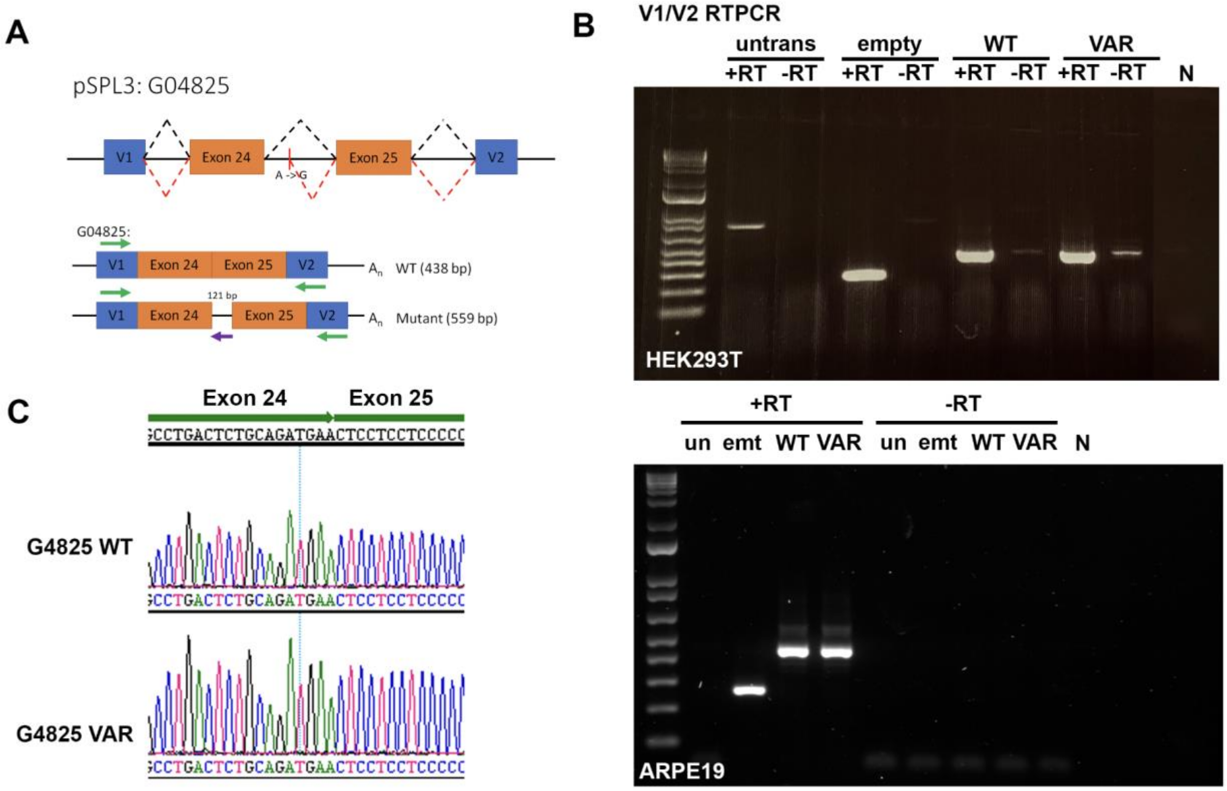


**Supplemental Figure 10. Predicted Splice Variant G04825 does not use alternative splice site in minigene assays.** Schematic of mini gene assay performed to determine effects of intronic variant on splicing and expected results (A). Variant c.3194+122A>G is spliced properly in HEK293T or ARPE-19 cells (B). Lanes included untransfected cells (un or untrans), empty vector, wild type (WT) MYRF exon 24 and 25, and the c.3194+122A>6 variant (VAR). Chromatograms from DNA sequencing the PCR products only detected normal splice site usage in mutants (C).

### **Supplemental Table 1.** Shared DEGs in Myrf$^{humdG/humdG}$ and RxCre;Myrf$^{fl/fl}$ RPE.

| Gene | Humanized Mouse Model (Myrf hWT/hMUT) | | | | | Conditional KO Model (RxCre;Myrf fl/fl) | | | | |
|---|---|---|---|---|---|---|---|---|---|---|
| | p_val.x | HumFC | pct.1.x | pct.2.x | p_val_adj.x | p_val.y | P0_FC | pct.1.y | pct.2.y | p_val_adj.y |
| Padi2 | 8.61E-09 | -2.4290381 | 0.038 | 0.132 | 2.10E-04 | 8.68E-08 | -1.9211933 | 0.016 | 0.138 | 0.00211807 |
| Ppp1r16b | 2.99E-17 | -2.184952 | 0.067 | 0.248 | 7.30E-13 | 3.49E-10 | -4.4825747 | 0.012 | 0.164 | 8.52E-06 |
| A930028N01 | 5.68E-12 | -1.6914712 | 0.12 | 0.274 | 1.39E-07 | 8.57E-25 | -6.477241 | 0.004 | 0.337 | 2.09E-20 |
| Rasgrp3 | 7.43E-20 | -1.5360456 | 0.195 | 0.427 | 1.81E-15 | 6.67E-27 | -5.7056312 | 0.039 | 0.401 | 1.63E-22 |
| Col5a2 | 9.16E-26 | -1.3483427 | 0.172 | 0.476 | 2.24E-21 | 4.96E-10 | -3.1773161 | 0.181 | 0.361 | 1.21E-05 |
| Tmem229b | 4.67E-11 | -1.3379554 | 0.134 | 0.285 | 1.14E-06 | 1.51E-08 | -3.3814417 | 0.094 | 0.25 | 3.69E-04 |
| Col3a1 | 4.43E-52 | -1.2899615 | 0.538 | 0.939 | 1.08E-47 | 7.13E-37 | -4.6805495 | 0.157 | 0.603 | 1.74E-32 |
| Dcn | 1.19E-28 | -1.2454641 | 0.258 | 0.588 | 2.91E-24 | 6.80E-19 | -2.6825776 | 0.217 | 0.514 | 1.66E-14 |
| Wfikkn2 | 5.61E-09 | -1.1103496 | 0.212 | 0.359 | 1.37E-04 | 2.17E-19 | -4.5431578 | 0.154 | 0.424 | 5.30E-15 |
| Col1a1 | 4.23E-48 | -1.108001 | 0.746 | 0.977 | 1.03E-43 | 1.04E-68 | -4.2314607 | 0.287 | 0.9 | 2.54E-64 |
| Col5a1 | 1.69E-16 | -1.1008552 | 0.162 | 0.386 | 4.11E-12 | 1.14E-11 | -4.1803179 | 0.146 | 0.342 | 2.78E-07 |
| Col1a2 | 2.14E-46 | -1.0730269 | 0.697 | 0.976 | 5.22E-42 | 8.60E-56 | -3.2611329 | 0.555 | 0.923 | 2.10E-51 |
| Cdh3 | 1.10E-10 | -1.0686517 | 0.309 | 0.479 | 2.69E-06 | 1.04E-06 | -1.4447087 | 0.547 | 0.534 | 0.02533827 |
| Ramp2 | 6.20E-10 | -1.0468211 | 0.177 | 0.328 | 1.51E-05 | 1.70E-36 | -5.2070951 | 0.031 | 0.492 | 4.16E-32 |
| Fbln2 | 2.51E-09 | -1.0078024 | 0.149 | 0.295 | 6.13E-05 | 1.60E-13 | -3.4955629 | 0.087 | 0.31 | 3.90E-09 |
| Upk1b | 1.65E-09 | -0.982261 | 0.135 | 0.275 | 4.02E-05 | 1.07E-24 | -5.8897906 | 0.016 | 0.352 | 2.62E-20 |
| Col6a3 | 8.51E-20 | -0.9553488 | 0.143 | 0.413 | 2.08E-15 | 3.00E-11 | -3.0709538 | 0.031 | 0.215 | 7.33E-07 |
| Col14a1 | 2.10E-16 | -0.9100164 | 0.139 | 0.375 | 5.14E-12 | 3.33E-08 | -5.074051 | 0.02 | 0.148 | 8.13E-04 |
| Upk3b | 7.60E-09 | -0.8336197 | 0.231 | 0.382 | 1.86E-04 | 8.86E-33 | -6.3087168 | 0.012 | 0.432 | 2.16E-28 |
| Col12a1 | 1.77E-10 | -0.813864 | 0.103 | 0.258 | 4.31E-06 | 4.63E-14 | -6.6053469 | 0.016 | 0.222 | 1.13E-09 |
| Gucy1a1 | 3.98E-08 | -0.7656024 | 0.156 | 0.292 | 9.71E-04 | 4.02E-07 | -2.6182813 | 0.118 | 0.262 | 0.00982239 |
| Lum | 7.71E-09 | -0.6763242 | 0.141 | 0.289 | 1.88E-04 | 5.24E-25 | -6.1675238 | 0.024 | 0.366 | 1.28E-20 |
| Sox10 | 2.51E-06 | -0.6366696 | 0.156 | 0.269 | 0.06130799 | 1.51E-15 | -2.7631164 | 0.035 | 0.275 | 3.69E-11 |
| Vim | 4.83E-18 | -0.6033612 | 0.655 | 0.923 | 1.18E-13 | 1.37E-08 | -0.5002305 | 0.878 | 0.84 | 3.35E-04 |
| Gm42418 | 4.43E-27 | -0.4386939 | 1 | 1 | 1.08E-22 | 1.25E-29 | -0.8836091 | 1 | 1 | 3.04E-25 |
| Timp3 | 5.68E-07 | -0.4129193 | 0.342 | 0.513 | 0.01385337 | 1.55E-06 | -1.085068 | 0.689 | 0.638 | 0.03771103 |
| Cpe | 2.71E-07 | -0.352342 | 0.33 | 0.511 | 0.00662045 | 1.23E-09 | -2.2066684 | 0.299 | 0.444 | 3.01E-05 |
| Hist1h4h | 1.48E-59 | 2.37574774 | 0.693 | 0.301 | 3.62E-55 | 7.73E-07 | 0.52701568 | 0.236 | 0.1 | 0.01887067 |
| Hist1h2ac | 2.51E-07 | 2.06143437 | 0.12 | 0.043 | 0.00611402 | 1.29E-08 | 1.1057587 | 0.193 | 0.06 | 3.15E-04 |
| Nnat | 3.55E-26 | 1.77823449 | 0.557 | 0.323 | 8.66E-22 | 7.42E-07 | 0.36941756 | 0.461 | 0.251 | 0.01811645 |
| Hist1h1d | 9.09E-20 | 1.5992939 | 0.496 | 0.295 | 2.22E-15 | 2.41E-07 | 0.94317392 | 0.154 | 0.046 | 0.00588459 |
| Id1 | 2.38E-09 | 1.24813453 | 0.365 | 0.236 | 5.82E-05 | 2.70E-23 | 1.38557221 | 0.701 | 0.332 | 6.60E-19 |
| Id3 | 2.37E-09 | 1.09360655 | 0.5 | 0.419 | 5.78E-05 | 5.48E-44 | 1.59423662 | 0.894 | 0.607 | 1.34E-39 |
| Csrp2 | 2.21E-06 | 1.03813275 | 0.416 | 0.332 | 0.05404357 | 4.56E-22 | 1.1569519 | 0.795 | 0.488 | 1.11E-17 |
| Ldha | 2.76E-30 | 0.91195372 | 0.828 | 0.718 | 6.74E-26 | 4.30E-21 | 0.83002459 | 0.878 | 0.679 | 1.05E-16 |
| Igfbp2 | 2.20E-12 | 0.85547762 | 0.622 | 0.5 | 5.38E-08 | 4.42E-23 | 1.33520614 | 0.87 | 0.719 | 1.08E-18 |
| Pgam1 | 1.99E-14 | 0.73640801 | 0.695 | 0.615 | 4.86E-10 | 3.93E-10 | 0.41017261 | 0.898 | 0.725 | 9.60E-06 |
| Tpi1 | 6.69E-13 | 0.67554877 | 0.716 | 0.645 | 1.63E-08 | 4.16E-11 | 0.54673703 | 0.894 | 0.703 | 1.01E-06 |
| Pkm | 6.61E-16 | 0.63555949 | 0.796 | 0.722 | 1.61E-11 | 1.74E-16 | 0.5262821 | 0.961 | 0.787 | 4.25E-12 |
| Eno1 | 1.29E-10 | 0.59223741 | 0.697 | 0.635 | 3.15E-06 | 1.73E-15 | 0.59974977 | 0.906 | 0.707 | 4.22E-11 |
| Pgk1 | 1.31E-06 | 0.53854368 | 0.662 | 0.614 | 0.03194289 | 5.71E-25 | 1.02859633 | 0.898 | 0.641 | 1.39E-20 |
| Rps29 | 1.36E-30 | 0.52578933 | 0.996 | 0.997 | 3.33E-26 | 3.31E-11 | 0.2963416 | 0.996 | 0.98 | 8.07E-07 |
| Rps13 | 4.86E-16 | 0.48302699 | 0.96 | 0.947 | 1.19E-11 | 3.18E-09 | 0.29802274 | 0.984 | 0.927 | 7.77E-05 |
| Rps10 | 6.03E-19 | 0.48209411 | 0.973 | 0.973 | 1.47E-14 | 1.98E-08 | 0.26850535 | 0.984 | 0.953 | 4.84E-04 |
| Rpl15 | 2.81E-15 | 0.46655307 | 0.935 | 0.916 | 6.85E-11 | 7.44E-11 | 0.31861905 | 0.949 | 0.914 | 1.82E-06 |
| Rps28 | 5.94E-17 | 0.45964916 | 0.954 | 0.957 | 1.45E-12 | 4.32E-08 | 0.27782396 | 0.941 | 0.913 | 0.00105487 |
| Atp5g2 | 1.15E-06 | 0.45963404 | 0.687 | 0.652 | 0.02817153 | 1.02E-13 | 0.46302427 | 0.909 | 0.78 | 2.49E-09 |
| Ppia | 1.22E-19 | 0.45182984 | 0.983 | 0.96 | 2.98E-15 | 5.28E-12 | 0.31265461 | 1 | 0.982 | 1.29E-07 |
| Gapdh | 6.77E-11 | 0.44384789 | 0.891 | 0.859 | 1.65E-06 | 3.61E-23 | 0.55224595 | 0.98 | 0.936 | 8.80E-19 |
| Rps18 | 6.93E-17 | 0.43336289 | 0.96 | 0.957 | 1.69E-12 | 4.06E-10 | 0.27620586 | 0.984 | 0.956 | 9.91E-06 |
| Eef1b2 | 2.87E-06 | 0.41823839 | 0.8 | 0.811 | 0.06994704 | 1.13E-07 | 0.33853738 | 0.925 | 0.821 | 0.00275352 |
| Tuba1b | 1.22E-06 | 0.40039428 | 0.731 | 0.751 | 0.0298906 | 2.05E-07 | 0.41470986 | 0.921 | 0.831 | 0.0050137 |
| Rps24 | 8.90E-18 | 0.39327435 | 0.998 | 0.996 | 2.17E-13 | 2.10E-08 | 0.27256875 | 0.988 | 0.967 | 5.12E-04 |
| Rpl29 | 9.54E-08 | 0.39008114 | 0.847 | 0.858 | 0.00232914 | 1.22E-08 | 0.29023579 | 0.961 | 0.883 | 2.98E-04 |
| Fau | 2.76E-13 | 0.36776227 | 0.981 | 0.977 | 6.74E-09 | 4.03E-11 | 0.27661819 | 0.984 | 0.949 | 9.84E-07 |
| Rpl41 | 1.17E-12 | 0.34505763 | 0.969 | 0.983 | 2.85E-08 | 1.13E-28 | 0.49267313 | 0.996 | 0.974 | 2.77E-24 |
| Rps7 | 8.81E-10 | 0.33816186 | 0.968 | 0.962 | 2.15E-05 | 8.48E-07 | 0.27563809 | 0.988 | 0.918 | 0.02069986 |
| Rpl11 | 1.37E-09 | 0.33690149 | 0.968 | 0.949 | 3.34E-05 | 1.15E-12 | 0.32537874 | 0.988 | 0.938 | 2.80E-08 |
| Rpl3 | 3.59E-08 | 0.32687341 | 0.895 | 0.92 | 8.76E-04 | 2.51E-06 | 0.26231742 | 0.957 | 0.911 | 0.06115496 |
| Rpl23 | 1.81E-13 | 0.32420861 | 0.989 | 0.996 | 4.42E-09 | 8.75E-10 | 0.2918244 | 0.992 | 0.96 | 2.14E-05 |
| Rpl19 | 4.75E-08 | 0.29635087 | 0.947 | 0.952 | 0.00115988 | 4.59E-11 | 0.30327259 | 0.992 | 0.936 | 1.12E-06 |
| Rps19 | 1.61E-06 | 0.27159438 | 0.971 | 0.981 | 0.03935115 | 1.84E-07 | 0.27631867 | 0.976 | 0.944 | 0.00448985 |
| Rpl26 | 3.61E-06 | 0.27092542 | 0.95 | 0.949 | 0.08816051 | 7.76E-11 | 0.29954988 | 0.969 | 0.923 | 1.89E-06 |

Gene had to have avg_log2FC >= abs(0.25) for BOTH genotypes
Gene had to have p_adj_val <= 0.1 for BOTH genotypes

### **Supplemental Table 2.** Clinical features of MYRF nanophthalmos families.

Supplemental Table 2: Clinical features of MYRF nanophthalmos families

| Patient # | Family # | Sex | Ethnicity | Age at exam | Sporadic/ Familial | logMAR | | Phakic Refraction (SE) | | Lens status | | Axial Length (mm) | | Phakic ACD | | C/D Ratio | | Max Clinic IOP | | Scleral Thickness | | Glaucoma | Narrow angle? | Retinal folds | Choroidal folds | Pigmentary Retinopathy | Complications |
|---|---|---|---|---|---|---|---|---|---|---|---|---|---|---|---|---|---|---|---|---|---|---|---|---|---|---|---|
| | | | | | | OD | OS | OD | OS | OD | OS | OD | OS | OD | OS | OD | OS | OD | OS | OD | OS | | | | | | |
| P04818 | F2 | F | EUWA | 53 | Familial | 0.0969 | 0.301 | NR | NR | Pseudo | Pseudo | 16.93 | 16.75 | 2.75 | 2.39 | 0.15 | 0.1 | 16.5 | 16 | 1.9 | 1.8 | + | + | - | + | - | aqueous misdirection |
| P04825 | F2 | F | EUWA | 75 | Familial | NR | NR | NR | NR | NR | NR | 18.77 | 18.67 | NR | NR | NR | NR | NR | NR | NR | NR | NR | NR | NR | NR | NR | NR |
| P01965 | F1 | F | EUWA | 60 | Sporadic | 0.3979 | NLP | 7.25 | 7.25 | Pseudo | Pros | 17.4 | 17.1 | NR | NR | 0.99 | ND | 38 | 40 | +thick | +thick | + | + | NR | NR | + | RD, phthisis OS |

### **Supplemental Table 3.** Splicing variants in silico analysis.

**Supplemental Table 3: Splicing variants in silico analysis**

| Family | Gene | Position (hg19) | cDNA change | SpliceAI score | Pangolin score | Expected effect | |
|---|---|---|---|---|---|---|---|
| F1 | *MYRF* | chr11:61536960 G>A | c.460+167G>A | 0.992 | 0.78 | AG | intron 4 pseudoexon insertion causing frameshift |
| F2 | *MYRF* | chr11:61551519 A>G | c.3194+122A>G | 0.8 | 0.49 | DG | Gain cryptic donor exon 25, or exon 24 skipping |
